# RTNet neural network exhibits the signatures of human perceptual decision making

**DOI:** 10.1101/2022.08.23.505015

**Authors:** Farshad Rafiei, Medha Shekhar, Dobromir Rahnev

## Abstract

Convolutional neural networks show promise as models of biological vision. However, their decision behavior, including the facts that they are deterministic and use equal number of computations for easy and difficult stimuli, differs markedly from human decision-making, thus limiting their applicability as models of human perceptual behavior. Here we develop a new neural network, RTNet, that generates stochastic decisions and human-like response time (RT) distributions. We further performed comprehensive tests that showed RTNet reproduces all foundational features of human accuracy, RT, and confidence and does so better than all current alternatives. To test RTNet’s ability to predict human behavior on novel images, we collected accuracy, RT, and confidence data from 60 human subjects performing a digit discrimination task. We found that the accuracy, RT, and confidence produced by RTNet for individual novel images correlated with the same quantities produced by human subjects. Critically, human subjects who were more similar to the average human performance were also found to be closer to RTNet’s predictions, suggesting that RTNet successfully captured average human behavior. Overall, RTNet is a promising model of human response times that exhibits the critical signatures of perceptual decision making.

## Introduction

Traditional cognitive models of perceptual decisions^1–4^ are able to account for the major features of human perceptual decision making, but do not operate on the level of images. Recently, convolutional neural networks (CNNs) have reached and sometimes exceeded human-level performance for novel images^5,6^. In addition, these networks naturally handle multi-choice categorization tasks and are promising models of the processing related to object recognition in the ventral visual stream of the human brain^5,7,8^. However, traditional CNNs’ decision behavior differs markedly from human decision behavior, thus limiting their applicability as models of human perceptual decision making. Specifically, unlike humans, traditional CNNs are both deterministic (i.e., they always give the same response for a given stimulus) and static (i.e., they are invariant in the amount of time spent on processing different images and thus always produce the same response time).

Several lines of work have tried to build mechanisms into neural networks to make them stochastic and dynamic^9–13^. Early research on shallow multi-layer perceptron models was able to create models that were both stochastic and dynamic. These models were able to explain human behavior on simple cognitive tasks^14–16^. However, these models are not image-computable (i.e., they cannot handle complex input such as images). More recent work has produced image-computable dynamic networks capable of generating response times (RTs) via mechanisms that allow computational resources utilized for the decision to increase with time^9–11^, thus allowing responses to evolve through each processing step. However, although these networks can mimic the speed-accuracy trade off (SAT) found in humans, they are deterministic and their internal mechanisms are not well supported by existing models of human perception and cognition. Finally, another class of models generates RTs using the biologically-inspired mechanism of recurrent processing^17–21^, which allows flexible modulation of a finite network’s computational power^10,22^. Nevertheless, these networks are also deterministic and have not been evaluated on the whole range of choice, RT, and confidence effects shown by humans.

Here we combine modern CNNs with traditional cognitive models to create a model that is image-computable, stochastic, and dynamic, and can reproduce the critical features of perceptual decision making for novel images. The model, which we call RTNet for its ability to model human RTs, features a deep convolutional neural network with noisy weights and processes a given image several times using a different random sample of these weights in each processing step (A). These weights are sampled from a Bayesian neural network (BNN) that estimates a posterior distribution over the best network parameters learnt during training. By sampling from these noisy weight distributions at each processing step, the network’s units produce variable responses to the same input that mimic the randomness of neural responses. After each processing step, RTNet accumulates the output corresponding to each choice until one of the choices reaches a predefined threshold. The model therefore has a strong conceptual relationship to race models from the cognitive literature on decision-making, which postulate a noisy accumulation process with separate accumulators for each choice^23–25^. By combining the image-computability of CNNs with traditional models of perception, we expect RTNet to be applicable across a wide range of perceptual tasks as well as reproduce the basic features of human perceptual decision making.

To assess a model’s ability to make decisions similar to humans, one needs to test whether it produces the foundational features of human decision-making^26^. Human perceptual decision making has been studied primarily in the context of 2-choice tasks using artificial stimuli such as Gabor patches or random dot motion^27^ (although notable exceptions exist where N-choice tasks are used^28–31)^. Therefore, we first replicate the known decision-making signatures from 2-choice tasks using an 8-choice task with meaningful images (hand-written digits taken from the MNIST dataset^32^). We manipulate 1) task difficulty by adding two different levels of noise to the images, and 2) speed-accuracy trade off (SAT) by asking subjects to emphasize either the accuracy or speed of their responses on different trials.

Critically, we test RTNet under the same conditions and with the same images seen by the human subjects to explore the model’s capability to produce behavior similar to human agents. Beyond testing whether RTNet can reproduce the basic features of human perceptual decision making, we also explore whether the accuracy, RT, and confidence produced by RTNet for individual images predict the corresponding quantities for humans on the same images. Finally, throughout the paper, we compare the behavior of RTNet to that of three other popular dynamic CNNs. The first model is Parallel Cascaded Network^9^ (CNet; Figure 1B), which is currently thought to be the best image-computable model that can mimic the SAT characteristics of humans^12^. The second is BLNet^10^, which belongs to a class of models that uses recurrent processing and has been validated on a range of perceptual tasks involving manipulations beyond SAT (Figure 1C). The third is Multi-Scale Dense Networks^13^ (MSDNet; D), which implements one of the most common ways for generating RTs in image-computable models. We find that RTNet’s behavior mimics human perceptual decision making better than all three of these CNNs.

**Figure 1.**
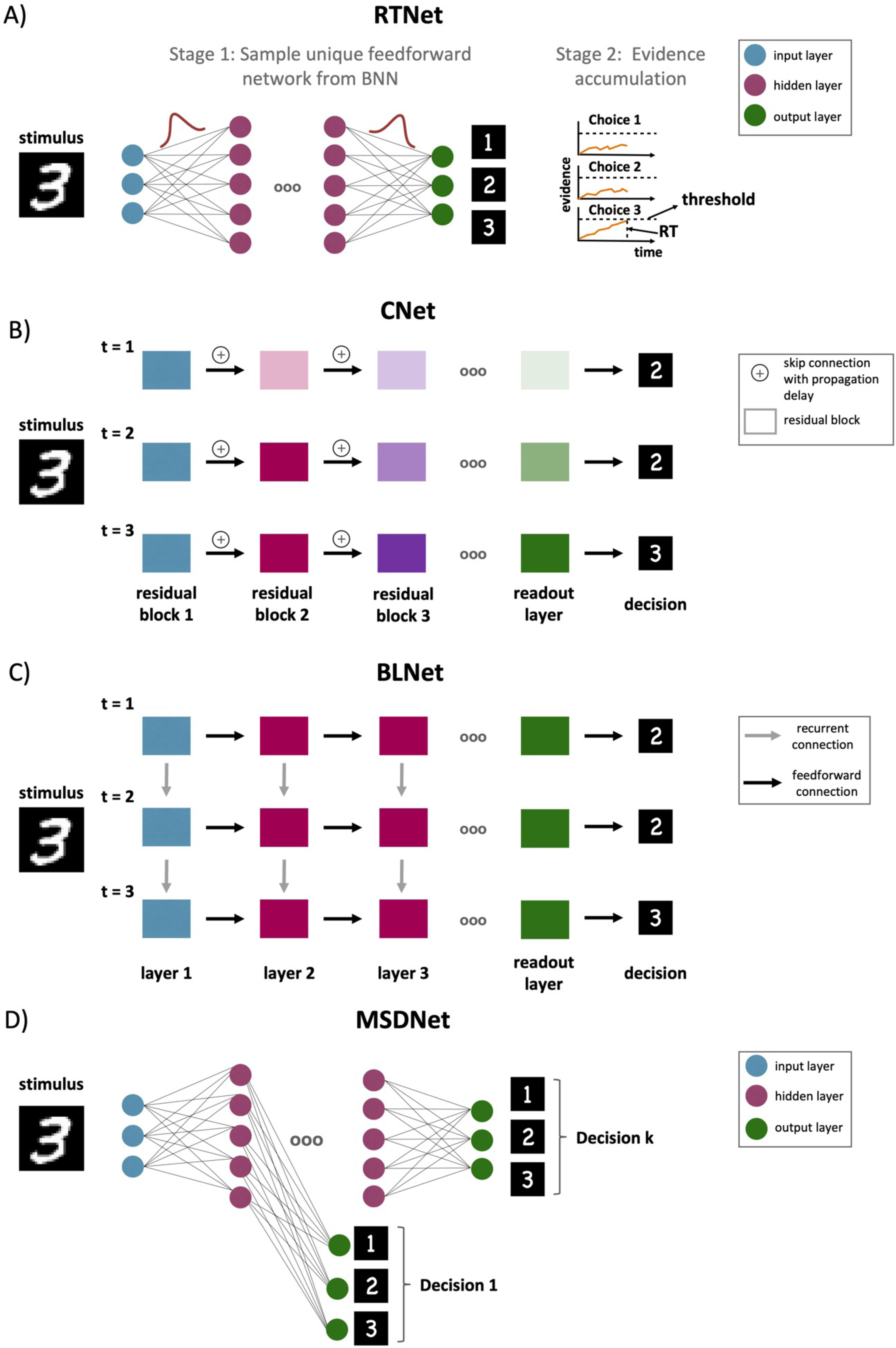
Model architectures. (A) RTNet architecture. Unlike standard CNNs, the connection weights in RTNet are not fixed but chosen from a distribution. A stimulus is processed multiple times by the network, each time using a different set of weights sampled randomly from a Bayesian neural network. The evidence from each processing step is accumulated and a decision is made when the evidence for one of the choices reaches a threshold. This architecture results in both stochastic decisions and variable RT. (B) Parallel cascaded network (CNet) architecture^9^. CNet introduce propagation delays between residual blocks (each of which consist of two convolutional layers). At each time step, all residual blocks parallelly receive inputs from lower blocks but due to propagation delays, earlier blocks achieve stable activations faster, whereas the later blocks require multiple processing steps to receive complete input and achieve stable activations. The network can generate a decision via the readout layer at any time step, although if the time step is less than the number of residual blocks, the decision will be based on partial input in later blocks. (C) BLNet architecture^10^. BLNet is a recurrent convolutional neural network (RCNN) with bottom-up and lateral recurrent connections. Time steps are defined in terms of the number of feedforward sweeps of the network. At each time-step, a layer receives feedforward input from the previous layer as well as recurrent input from its own activations at the previous time-step. The readout can be evaluated at each time-step to generate a response if it exceeds the threshold. The network’s can trade off speed and accuracy as higher thresholds require more feedforward and recurrent computations, effectively resulting in a deeper network being unrolled across time. (D) Multi-scale dense network (MSDNet) architecture^13^. In this network, each hidden layer features its own classifier allowing MSDNet to make a separate decision after the processing in each layer is completed. This allows the network to stop processing an image early if that image can already be decoded from earlier layers of the network, thus resulting in different RTs for different images.

## Results

We collected data from 60 human subjects who performed a digit discrimination task (**Error! Reference source not found.**A). The experiment was a 2 x 2 design with factors of task difficulty (easy vs. difficult images) and speed pressure (speed vs. accuracy focus). Each condition consisted of 120 unique images, and each subject made a decision regarding each image exactly twice, which allowed us to determine the level of stochasticity in human behavior (**Error! Reference source not found.**B). Overall, each subject completed 960 trials in total.

Having obtained these human data, we compared the human behavior to that of RTNet, CNet, BLNet, and MSDNet. Both RTNet and MSDNet were implemented using the eight-layer AlexNet architecture with five convolutional layers followed by three fully connected layers^33^. CNet was based on the architecture of ResNet18 since the implementation of this model relies on residual blocks and skip connections. Finally, for BLNet, we used the original architecture implemented by Spoerer et al.^10^, which consists of seven convolutional layers and a fully connected readout layer. Given that humans and deep learning models are impacted differently by stimulus noise^34,35^, we adjusted the noise levels of the images seen by each network to match their overall accuracy to the accuracy produced by the human subjects. In addition, to allow the networks to reproduce the speed-accuracy trade off observed in the human data, we adjusted the threshold value that triggers a decision for each CNN as to match the human accuracy separately in the speed-and accuracy-focused conditions. To improve the correspondence between the model predictions and the human data, we trained 60 instances of each model (by only changing the random initialization before training began) and analyzed the data produced by these 60 instances in equivalent manner to the 60 human subjects.

### Signatures of human perceptual decision making

We examined six foundational signatures of human perceptual decision making that have already been established in studies of 2-choice tasks: 1) Human decisions are stochastic, meaning that the same stimulus can elicit different responses on different trials^36,37^, 2) increasing speed stress shortens RT but decreases accuracy (speed-accuracy trade off)^26,38,39^, 3) more difficult decisions lead to reduced accuracy and longer RT^26,40,41^, 4) RT distributions are right-skewed, and this skew increases with task difficulty^26^, 5) RT is lower for correct than for error trials^41–45^, and 6) confidence is higher for correct than for error trials^46^. For each of these signatures, we confirmed that the signature also occurs for our 8-choice task with naturalistic images, and then tested whether RTNet, CNet, BLNet and MSDNet exhibit the same signature.

#### Stochasticity of human decisions

A central feature of human behavior is that human decisions are stochastic such that the same stimulus can elicit different responses on different trials^36,37,47^. We quantified the level of stochasticity in each condition by presenting each image twice. We first confirmed that our estimates of human stochasticity were robust and reliable by showing that similar estimates are obtained when analyzing the odd vs. even numbered subjects (Supplementary Figure 1). On average across all conditions, 36% of all images received different responses on the two presentations. A one-sided Wilcoxon signed-rank test showed that this observed frequency of stochastic responses is indeed significantly greater than zero (Z(59) = 32896, p < 0.001, rank-biserial correlation (effect size) = 1) (**Error! Reference source not found.**A). A repeated measures ANOVA with factors stimulus difficulty (easy vs. difficult) and SAT (speed vs. accuracy stress) revealed that stochasticity increased with both higher task difficulty (F(1,63) = 871.869, p < 0.001, 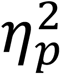 = 0.933) and higher speed pressure (F(1,63) = 9.135, p = 0.004, 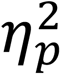 = 0.127).

Since RTNet uses a random sample of weights for each processing step, it naturally produces stochastic decisions too. On average across all conditions, RTNet produced different responses on the two image presentations on 20% of trials (One-sided Wilcoxon signed-rank test: Z(59) = 2892, p < 0.001, rank-biserial correlation (effect size) = 1; **Error! Reference source not found.**B). This level of stochasticity was lower than for human subjects and stems from the fact that the variability in the weights was fixed a priori by training a Bayesian neural network. However, it is possible for RTNet to match the levels of stochasticity observed in humans by increasing the variability of the network’s weights. Indeed, we confirmed that the stochasticity of the decisions made by RTNet can be robustly manipulated by changing the variability of its weight distributions (Supplementary Figure 2). Further, the stochasticity in human decisions partially stems from factors such as fluctuations in attention, arousal, or serial dependence^36,37,47,48^, which we did not attempt to model. Because of these considerations, we did not try to match RTNet to the exact level of human decision stochasticity observed in the data. Critically, however, RTNet exhibited the same features such that stochasticity increased with higher task difficulty (F(1,59) = 120.124, p < 0.001, 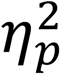 = 0.671) and higher speed stress (F(1,59) = 87.730, p < 0.001, 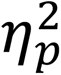 = 0.598).

On the other hand, for a fixed level of speed-accuracy trade off, CNet, BLNet and MSDNet are fully deterministic and do not exhibit any decision stochasticity (**Error! Reference source not found.**C-E). We note that it should be possible to add noise in the weights of these models to induce stochastic decisions, but such noise would decrease their accuracy much more than it affects RTNet given that only RTNet is able to average out the noise over repeated processing steps. Because RTNet is the only model that incorporates a mechanism for generating stochastic responses, these stochasticity analyses a priori favor RTNet over the other models. However, the rest of our analyses compare the behavior and predictions of models across a range of stimulus manipulations in which no model is a priori expected to be favored over the others.

#### Speed-accuracy trade off

The ability to trade off speed and accuracy against each other is a hallmark of decision-making across humans and many other animal species^38,39^. The human data confirmed that increased speed pressure led to lower accuracy (F(1,59) = 4.274, p = 0.043, 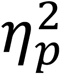 = 0.068; **Error! Reference source not found.**A) and shorter RTs (F(1,59) = 119.29, p < 0.001, 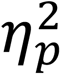 = 0.964; **Error! Reference source not found.**B). We also found a significant interaction between SAT and task difficulty for accuracy such that the SAT effect was greater for easy images (F(1,59) = 5.71, *p* = 0.020, 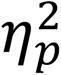 = 0.088). For RTs, however, we observed the opposite pattern where the SAT effect was heightened for difficult images (F(1,59) = 22.423, *p* < 0.001, 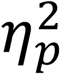 = 0.275). These results replicate findings from a previous study examining the effects of SAT manipulations on accuracy and RT as a function of stimulus contrast^49^. Further, as shown before^49^, these findings are also in line with predictions of the drift diffusion model (DDM), which is currently the standard model for explaining human RTs^1,2^.

All models were able to replicate the speed-accuracy trade off observed in humans. Increased speed pressure resulted in lower accuracy for RTNet (F(1,59) = 9.683, p = 0.003, 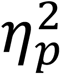 = 0.141), CNet (F(1,59) = 50.025, p < 0.001, 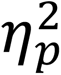 = 0.459), BLNet (F(1,59) = 11.611, p = 0.001, 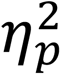 = 0.164), and MSDNet (F(1,59) = 21.841, p < 0.001, 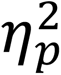 = 0.270). Increased speed pressure also led to shorter RTs for RTNet (F(1,59) = 3362.567, p < 0.001, 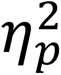 = 0.983), CNet (F(1,59) = 695.878, p < 0.001, 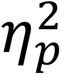 = 0.922), BLNet (F(1,59) = 607.093, p < 0.001, 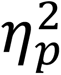 = 0.911), and MSDNet (F(1,59) = 584.081, p < 0.001, 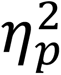 = 0.908). We note that the SAT manipulation had a relatively small effect on accuracy (1.04% for easy and 1.24% for difficult conditions for RTNet; the effects for the rest of the networks were of similar magnitude; Figure 4A). However, despite the small effect size, these effects were generally consistent across the 60 model instances (for RTNet, 54/60 instances showed the effect for easy images and 42/60 showed the effect for difficult images).

**Figure 2.**
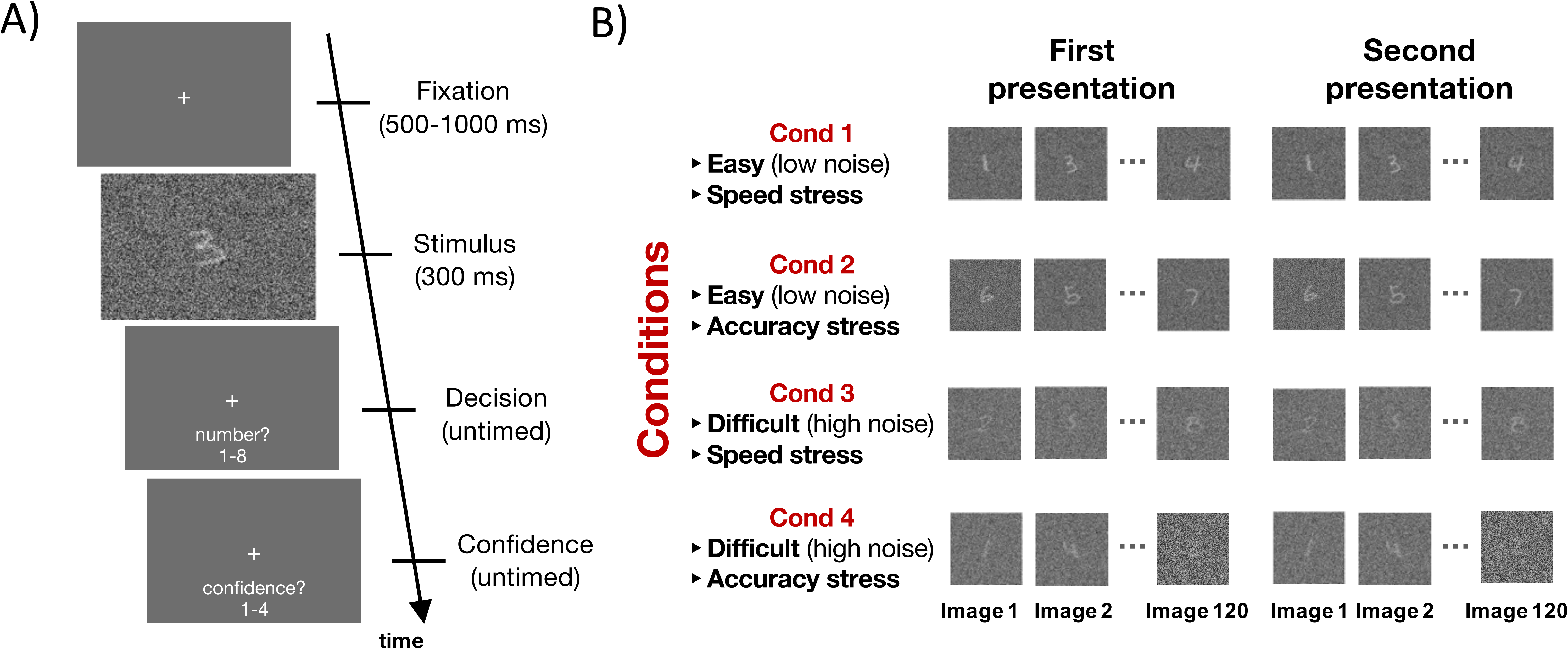
Experiment task. (A) Trial structure. Each trial began with a fixation cross presented for 500 to 1000 ms, followed by an image of a hand-written digit from the MNIST dataset embedded in noise and presented for 300 ms. Only the digits 1-8 were used. Subjects reported their choice and confidence (on a 4-point scale) using separate, untimed button presses. Note that the noisy stimulus subtended a visual angle of 6.06° and did not cover the entire screen. (B) Experimental design. The experiment included four conditions such that subjects judged easy (low noise) or difficult (high noise) images while emphasizing either speed or accuracy. Each condition featured 120 unique images that were the same across all subjects (total of 480 unique images in the experiment). In addition, each image was presented twice to allow the estimation of the stochasticity of human perceptual choices. Each subject thus completed a total of 960 trials. The images within the first and second sets of presentation were shown in a different random order.

**Figure 3.**
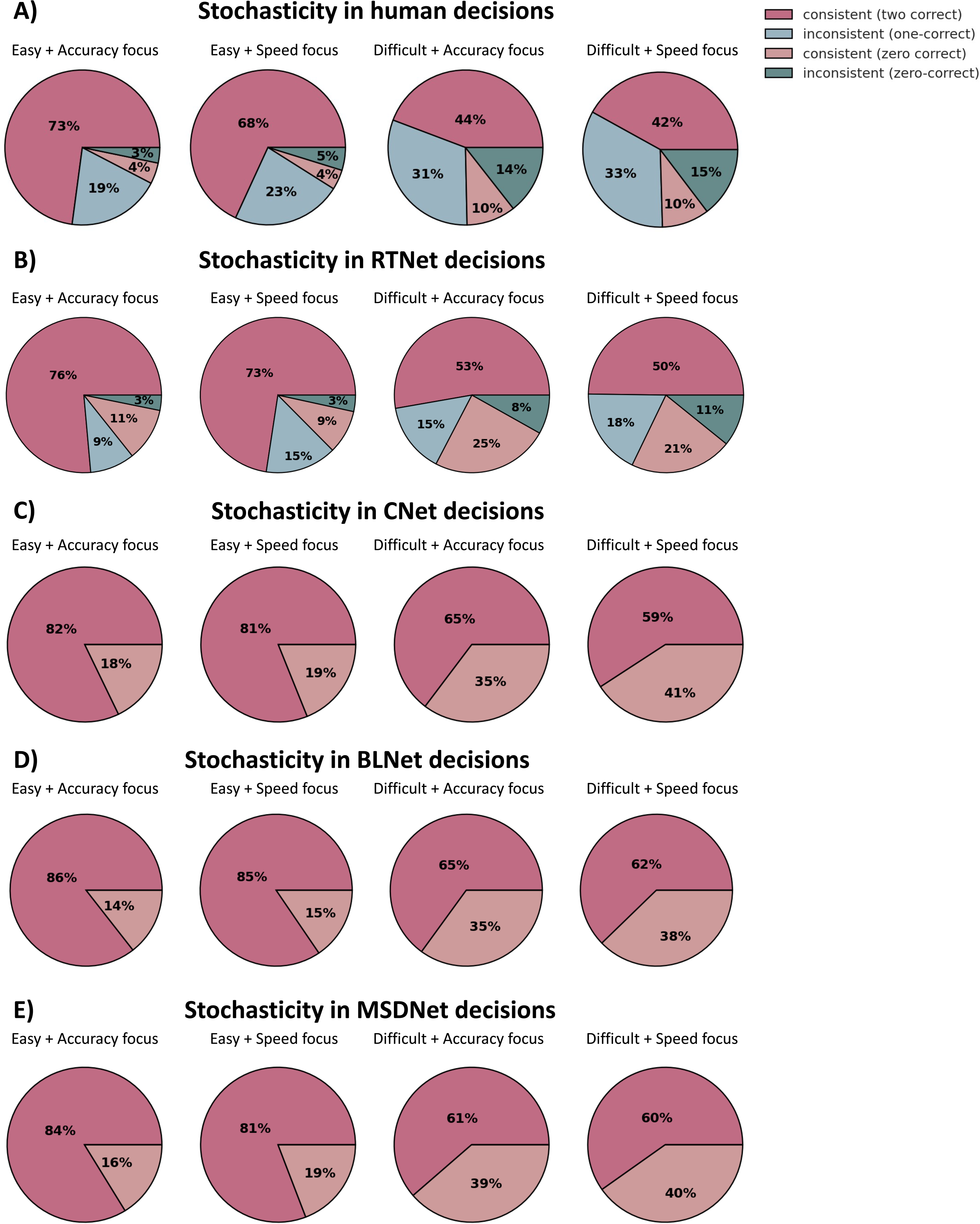
Decision stochasticity in humans and all networks. Stochasticity of decisions made by (A) humans, (B) RTNet, (C) CNet (D) BLNet, and (E) MSDNet. Warm colors indicate that the same response was given both times an image was presented (whether the response was correct or incorrect), whereas cool colors indicate that different responses were given for the two image presentations (whether or not any of them was correct). Humans and RTNet exhibit stochastic decision-making with stochasticity increasing with task difficulty and speed stress. However, CNet, BLNet and MSDNet in their standard versions are fully deterministic. In the legend, “consistent (two correct)” refers to instances when the correct responses was given for both presentations of a given image, “consistent (zero correct)” refers to instances when the same incorrect choice was made both times, “inconsistent (one correct)” refers to instances when only one of the choices was correct, “inconsistent (zero correct)” refers to instances where different incorrect choices were made each time.

**Figure 4.**
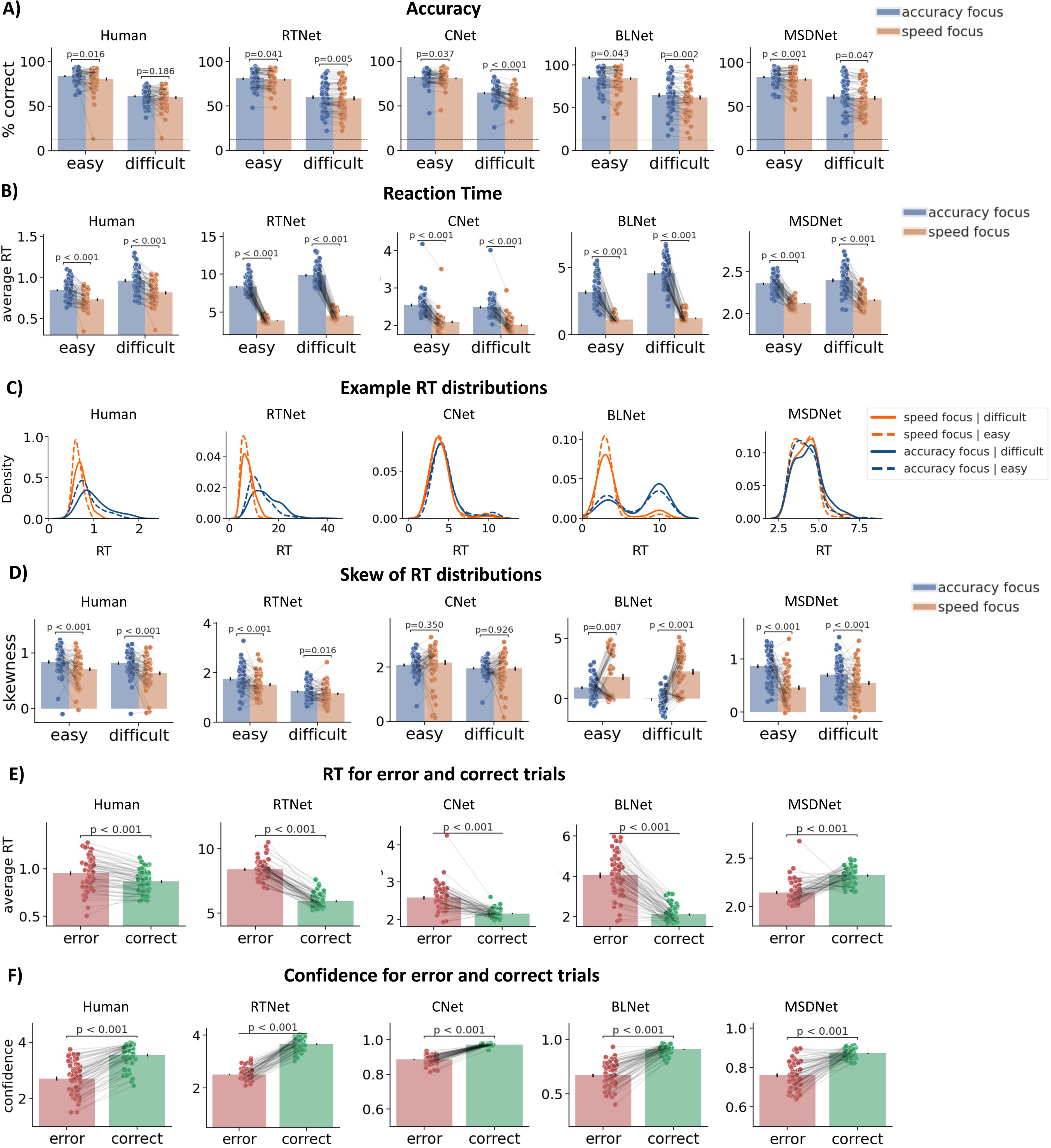
Behavioral effects shown by human subjects and the models. (A) Accuracy for humans (n = 60) decreases when response speed is emphasized as well as for more difficult decisions. Both effects are exhibited by all the networks (n = 60 model instances). (B) RT for humans becomes shorter when response speed is emphasized, as well as for easier decisions. Both effects are also exhibited robustly by RTNet and BLNet. However, while both CNet and MSDNet produced a robust effect for the speed manipulation, they exhibited much smaller effects for the difficulty manipulation. RT for humans is measured in seconds and RT for the networks is measured in the number of steps over which evidence is accumulated (for RTNet), propagation steps (for CNet), feedforwards sweeps (for BLNet) and number of layers (for MSDNet). (C) RT distributions for a representative subject/model. (D) The skewness of RT distributions change across conditions. For humans and RTNet, the skewness of the RT distributions was higher for easier tasks and for accuracy focus. However, CNet, BLNet, and MSDNet showed clear deviations from the human pattern of results. (E) For humans, RTNet, CNet and BLNet, two-sided paired t-tests showed that error trials were associated with higher RT than correct trials. However, MSDNet showed the opposite pattern such that correct trials were associated with longer processing time. (F) Confidence for correct trials was higher than confidence for error trials for humans and all networks. For all panels, dots represent individual subjects; error bars show SEM. The p-values are derived from two-sided Wilcoxon’s signed rank tests (for mean RT comparisons) and two-sided paired t-tests (for all other measures).

The SAT manipulation had a much stronger effect on RTs compared to accuracy, which may be attributed to the fact that RTs are a more sensitive measure of performance. Further, the SAT effect on RTs was much stronger for humans, RTNet and BLNet, compared to the other models. The individual RT distributions show a clear separation between the speed and accuracy focus conditions for humans, RTNet and BLNet but not for CNet and MSDNet (Figure 4C). Nevertheless, these results indicate that speed-accuracy trade off is robustly observed even for relatively complex task with naturalistic images, and that all models examined here exhibit this foundational phenomenon.

#### Difficult decisions lead to reduced accuracy and longer RT

Another ubiquitous feature of decision-making is that more difficult stimuli lead to lower accuracy and longer RT^26,50^. Our human data robustly showed this effect with more difficult stimuli leading to lower accuracy (F(1,59) = 1558.500, *p* < 0.001, 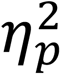 = 0.964; **Error! Reference source not found.**A) and longer RT (F(1,59) = 411.154, *p* < 0.001, 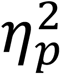 = 0.875; **Error! Reference source not found.**B). The same pattern was robustly observed for RTNet and BLNet, where difficult stimuli led to lower accuracy (RTNet: F(1,59) = 218.510, *p* < 0.001, 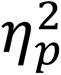 = 0.787; BLNet: F(1,59) = 200.543, *p* < 0.001, 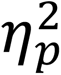 = 0.773) but longer RT (RTNet: F(1,59) = 233.452, *p* < 0.0001, 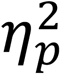 = 0.798; BLNet: F(1,59) = 186.604, *p* < 0.001, 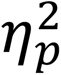 = 0.760). However, while CNet and MSDNet also showed a very robust effect on accuracy (CNet: F(1,59) = 1116.800, *p* < 0.001, 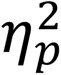 = 0.950; MSDNet: F(1,59) = 247.520, *p* < 0.001, 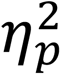 = 0.808), they exhibited a smaller effect for RT (CNet: F(1,59) = 11.070, *p* = 0.016, 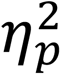 = 0.158; MSDNet: F(1,59) = 6.171, *p* = 0.002, 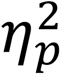 = 0.095). Indeed, out of the 60 model instances, only 23 CNet instances and 36 MSDNet instances exhibited an RT increase for more difficult stimuli, while this effect was present in 60/60 human subjects, 58/60 RTNet instances, and 59/60 BLNet instances. These results indicate that the effect of task difficulty on accuracy is exhibited robustly in humans and all networks, but the effect of task difficulty on RT is larger for humans, RTNet and BLNet compared to CNet and MSDNet (see Discussion).

#### Skewness of RT distributions

For simple 2-choice decisions, human RT distributions are generally positively skewed and the skewness changes as a function of task conditions^2,26^. Our 8-choice task produced RT distributions that closely resemble what is observed in standard 2-choice tasks (**Error! Reference source not found.**C). Similar-looking RT distributions were produced by RTNet but CNet and MSDNet produced RT distributions that, while still right-skewed, exhibited qualitative differences in their shapes (**Error! Reference source not found.**C). BLNet, on the other hand, produced RT distributions that were frequently bimodal and left-skewed.

We further assessed how the skewness of the RT distributions changed under different conditions. In humans, we found higher skewness for accuracy compared to speed focus (F(1,59) = 32.837, *p* < 0.001, 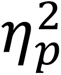 = 0.358), as well as for easy compared to difficult stimuli (F(1,59) = 5.098, *p* = 0.028, 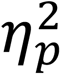 = 0.080; **Error! Reference source not found.**D). RTNet exhibited the same pattern with skewness increasing with a focus on accuracy (F(1,59) = 19.077, p < 0.001, 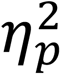 = 0.244) and with easier stimuli (F(1,59) = 93.342, p < 0.001, 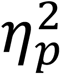 = 0.613). For CNet, we found no difference in skewness of RT distributions between the SAT conditions (F(1,59) = 0.428, p = 0.515, 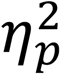 = 0.007), but skewness increased for easy compared to difficult stimuli (F(1,59) = 8.612, p = 0.005, 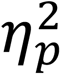 = 0.127). BLNet showed the opposite pattern to CNet with skewness increasing for the speed-focus condition (F(1,59) = 39.219, p < 0.001, 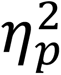 = 0.399) and failed to show difference in skewness between the easy and difficult stimuli (F(1,59) = 3.517, p = 0.066, 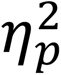 = 0.056). Finally, while MSDNet showed an increase in skewness with a focus on accuracy (F(1,59) = 64.866, *p* < 0.001, 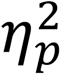 = 0.524), it produced RT distributions that did not significantly differ in skewness between the task difficulty conditions (F(1,59) = 1.259, *p* = 0.266, 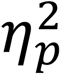 = 0.021). Overall, RTNet produced RT distributions which reflected the observed patterns in human data better than all other networks. It should be noted that CNet, BLNet, and MSDNet can only produce distinct RTs that are less than or equal to their layer numbers or residual blocks, which may affect their ability to reproduce human RT distributions unless a relatively high number of layers is used. On the other hand, RTNet can go through arbitrary number of samples regardless of the number of layers in its architecture.

#### RT is faster for correct compared to error trials

Another ubiquitous feature of human behavior in 2-choice tasks is that correct decisions are typically accompanied by faster RTs than incorrect decisions^41–45^. We replicated this effect in our 8-choice task (F(1,59) = 82.080, *p* < 0.001, 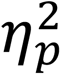 = 0.582; **Error! Reference source not found.**E). The same difference between correct and error RTs also emerged for RTNet (F(1,59) = 831.153, *p* < 0.001, 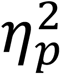 = 0.934), CNet (F(1,59) = 83.921, *p* < 0.001, 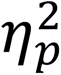 = 0.587), and BLNet (F(1,59) = 286.157, *p* < 0.001, 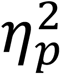 = 0.582). However, MSDNet exhibited the opposite pattern such that RTs were faster for error compared to correct trials (F(1,59) = 65.696, *p* < 0.001, 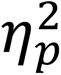 = 0.527). This behavior is due to the fact that errors produced by MSDNet come mostly from decisions made in earlier layers. It may be possible to reverse this behavior by using a much more conservative decision threshold in the early compared to the late layers of MSDNet, though the effectiveness of this strategy and its effect on all other behavioral signatures examined here would need to be tested. What is clear is that MSDNet in its current form makes a qualitatively wrong prediction regarding the difference between correct and error RT, whereas RTNet, CNet and BLNet naturally reproduce the empirical effect.

#### Confidence is higher for correct than error trials

Finally, a ubiquitous feature of confidence ratings is that they are higher for correct compared to incorrect decisions^46,51^. Our human data replicated this effect (F(1,59) = 472.172, *p* < 0.001, 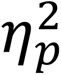 = 0.889; **Error! Reference source not found.**F). The effect was also robustly exhibited by all networks: RTNet (F(1,59) = 966.796, *p* < 0.001, 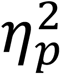 = 0.942), CNet (F(1,59) = 785.992, *p* < 0.001, 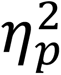 = 0.930), BLNet (F(1,59) = 374.031, *p* < 0.001, 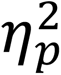 = 0.864), and MSDNet (F(1,59) = 131.923, *p* < 0.001, 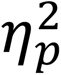 = 0.691). Therefore, humans and all networks robustly showed higher confidence for correct trials compared to incorrect trials.

### Model predictions of responses for individual images

The results above demonstrate that RTNet naturally reproduces all foundational features of human decision-making. On the other hand, CNet, BLNet, and MSDNet fail to exhibit stochastic decisions and skewness difference in RT distributions between the SAT/difficulty conditions, and MSDNet further fails to account for lower RT for correct decisions. However, RTNet’s ability in those respects can easily be matched by traditional cognitive models that do not work on image-level data^24,42,52^. Therefore, a critical advantage of RTNet over traditional cognitive models would be the ability to predict human behavior for individual, unseen images because traditional models cannot do that. Here we tested specifically whether the accuracy, RT, and confidence for unseen images produced by the networks predict the same quantities in humans.

#### Model predictions for individual subjects

In a first set of analyses, we assessed the correlations between the accuracy, RT, and confidence for each human subject and the corresponding quantities predicted by RTNet, CNet, BLNet, and MSDNet across all four conditions (easy with speed stress, difficult with speed stress, easy with accuracy stress, difficult with accuracy stress). We compared how well data from individual human subjects could be predicted by each model as well as from the data from the 59 remaining human subjects. This last quantity, which we call subject-to-group relationship, provides an estimate of the noise ceiling (i.e., the performance that a true model could achieve given inter-subject variability)^53^.

We found that all models predicted individual human data much better than chance for accuracy, RT, and confidence (two-sided one-sample t-tests, all *p*’s < 0.001, all Cohen’s *d* > 1.20). The one exception was BLNet, which had a weak negative correlation with human image-by-image accuracy (average *r* = −0.06, *p* = 0.002, Cohen’s *d* = 0.410, 95% CI = [-0.09, −0.02]). Critically, RTNet provided substantially better predictions than all other models (**Error! Reference source not found.**). Specifically, two-sided paired t-tests showed that RTNet produced better image-by-image predictions about accuracy (RTNet vs. CNet: t(59) = 30.672, *p* < 0.001, Cohen’s *d* = 4.747, 95% CI = [0.24, 0.27]; RTNet vs. BLNet: t(59) = 20.842, *p* < 0.001, Cohen’s *d* = 3.864, 95% CI = [0.37, 0.44]; RTNet vs. MSDNet: t(59) = 30.672, *p* < 0.001, Cohen’s *d* = 4.747, 95% CI = [0.24, 0.27]), RT (RTNet vs. CNet: t(59) = 18.638, *p* < 0.001, Cohen’s *d* = 2.370, 95% CI = [0.29, 0.35]; RTNet vs. BLNet: t(59) = 13.135, Cohen’s *d* = 0.472, 95% CI = [0.06, 0.08], *p* < 0.001; RTNet vs. MSDNet: t(59) = 13.318, *p* < 0.001, Cohen’s *d* = 1.152, 95% CI = [0.13, 0.18]), and confidence (RTNet vs. CNet: t(59) = 8.394, *p* < 0.001, Cohen’s *d* = 0.936, 95% CI = [0.07, 0.11]; RTNet vs. BLNet: t(59) = 6.587, *p* < 0.001, Cohen’s *d* = 0.391, 95% CI = [0.03, 0.05]; RTNet vs. MSDNet: t(59) = 7.68, *p* < 0.001, Cohen’s *d* = 0.471, 95% CI = [0.04, 0.06]).

Critically, RTNet’s predictions were reasonably close to the noise ceiling in all cases (calculated as the average subject-to-group correlation in the human data). Specifically, RTNet’s predictions were within 62.5%, 79.6%, and 64.8% of the noise ceiling for accuracy, RT, and confidence, respectively. These numbers were substantially lower for CNet (16.1%, 20.3%, 40.5%, respectively), BLNet (0%, 64.4%, 54.1%, respectively), and MSDNet (16.1%, 50%, and 51.3%, respectively). Thus, by reaching to between 62.5% and 79.6% of the noise ceiling, RTNet can provide excellent predictions for the accuracy, RT, and confidence produced by human subjects for images that the model was not trained on. Additionally, we derived the model predictions for averages across the 60 subjects across all conditions (Supplementary Figure 3) and found that RTNet still predicts average human accuracy and RT better than the other networks.

#### Model predictions within each condition separately

The analyses above explored the correlations between model predictions and human behavior across all experimental conditions. Because different conditions vary in their average accuracy, RT, and confidence, analyses across conditions are likely to produce higher correlations than if the same analyses are to be performed within each condition separately. Therefore, we repeated the analyses above but within each of the four conditions separately to investigate if the models can still account for accuracy, RT, and confidence on individual images. We found that RTNet, BLNet and MSDNet produced accuracy, RT, and confidence predictions that significantly correlate with individual subject data in all conditions (two-sided one-sample t-tests, all *p*’s < 0.001; Figure 6). However, while CNet produced accuracy and confidence predictions that significantly correlated with individual subject data in all conditions, its RT predictions for all conditions except accuracy focus with difficult images, were either zero or negative (p’s > 0.62).

**Figure 5.**
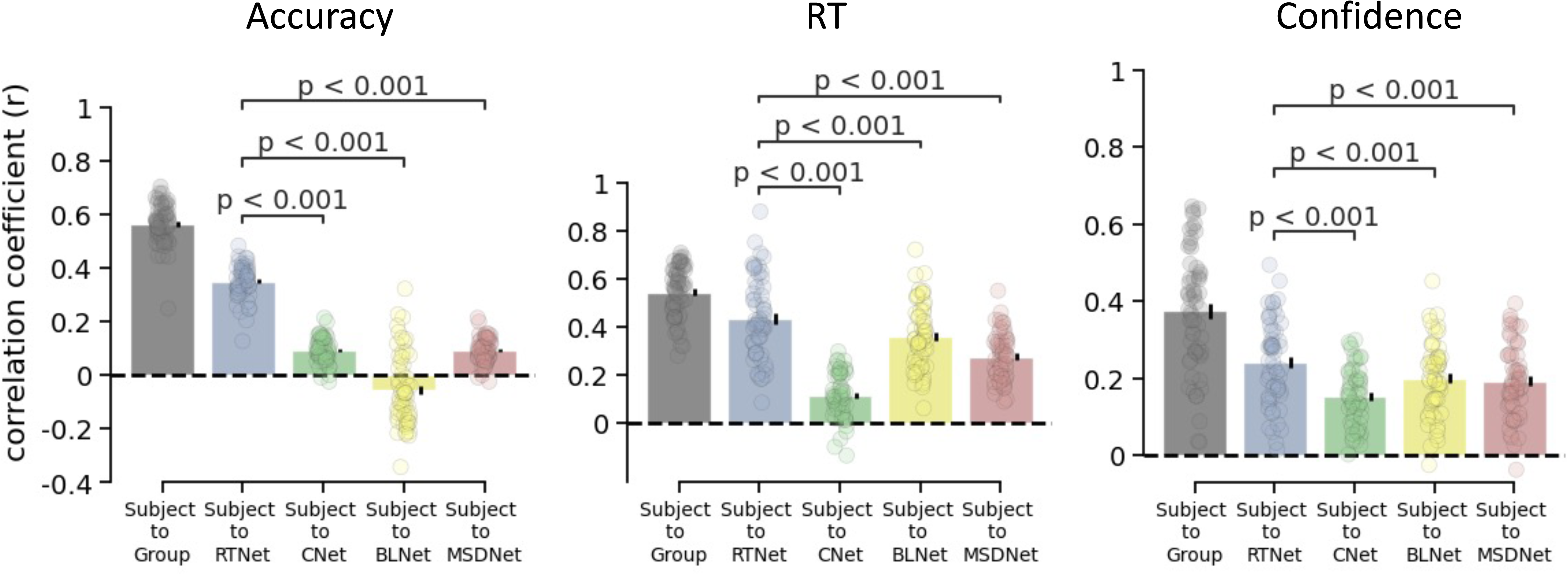
Image-by-image correlation between human data and each model across all experimental conditions for individual subjects. Correlation between data from individual human subjects (n = 60) and the group average, and correlations between data from individual subjects and the average of all 60 instances for RTNet, CNet, BLNet, and MSDNet. The correlations are computed separately for accuracy, RT, and confidence across all conditions. Critically, the correlation is stronger for RTNet than CNet, BLNet or MSDNet for each measure. The subject-to-group correlation provides an estimate of the noise ceiling for the network correlations. Dots represent individual subjects; error bars show SEM. The p-values are derived from two-sided paired t-tests.

**Figure 6.**
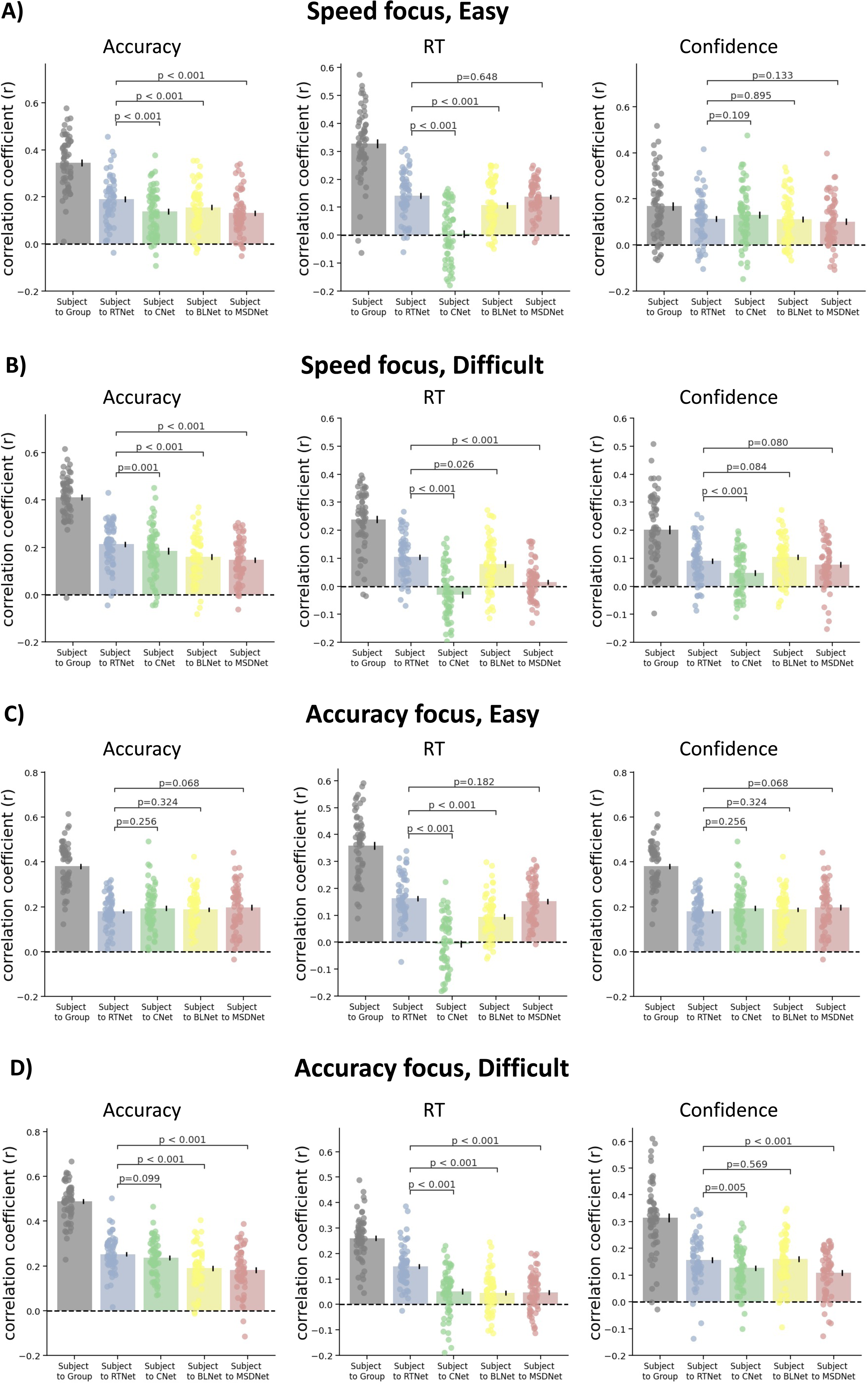
Image-by-image correlation between human data and each network within each experimental condition. Correlation between data from individual human subjects (n = 60) and the group average, as well as the average of all 60 instances for RTNet, CNet, BLNet, and MSDNet. The correlations are computed separately for accuracy, RT, and confidence within each experimental condition: A) speed focus; easy, B) speed focus; difficult, C) accuracy focus; easy, D) accuracy focus; difficult. The correlation is significantly stronger for RTNet compared to CNet (8/12 comparisons), BLNet (7/12 comparisons), and MSDNet (7/12 comparisons). RTNet never exhibits significantly weaker correlations than CNet, BLNet, or MSDNet. For all panels, dots represent individual subjects; error bars show SEM. The p-values are derived from two-sided paired t-tests.

Critically, however, RTNet predicted the individual data significantly better than the rest of the networks. Specifically, two-sided paired t-tests showed that RTNet provided better predictions than CNet in two out of four conditions for accuracy (all *p*’s < 0.001), in all four conditions for RT (all *p*’s < 0.0001), and in two out of four conditions for confidence (*p* < 0.005). Compared to BLNet, RTNet predicted individual data significantly better in three out of four conditions for accuracy (all *p*’s < 0.0001) and in all four conditions for RT (all *p*’s < 0.025). Compared to MSDNet, RTNet predicted the individual data significantly better in three out of four conditions for accuracy (all three *p*’s < 0.001) and in all four conditions for RT (all *p*’s < 0. 02). There was no significant difference in confidence predictions between RTNet and BLNet or between RTNet and MSDNet for any of the four conditions (all *p*’s > 0.05). RTNet was never significantly worse than CNet, BLNet or MSDNet in predicting any of the 12 comparisons. Overall, these results demonstrate that RTNet predicts human behavior well across all three measures and across different types of analyses (across-or within-condition), and does so better than CNet, BLNet and MSDNet.

### Humans more similar to the group are more similar to RTNet

Our subject-to-group analyses revealed substantial variability in how well individual subjects’ data corresponded to the group average (see Figure 5). Since the group average constitutes the best model of human behavior, one would expect that any good, generalizable model of behavior would also be able to capture this relationship between individual subjects and the group average. In other words, the strength of the relationship for an individual subject and the group should be linked to the strength of the relationship of that same subject and the model. Here we tested if such dependency holds true for RTNet, CNet, BLNet and MSDNet. We found that subjects who exhibited greater correlation in image-by-image accuracy across all conditions with rest of the group also exhibited greater correlation with the RTNet predictions (Pearson’s r = 0.685, *p* < 0.001, 95% CI = [0.52, 0.80]; Figure 7A). The same correspondence also emerged for RT (Pearson’s r = 0.825, *p* < 0.001, 95% CI = [0.72, 0.89]) and confidence (Pearson’s r = 0.894, *p* < 0.001, 95% CI = [0.83, 0.94]). Similar results were obtained for CNet (Accuracy: Pearson’s r = 0.389, *p* = 0.002, 95% CI = [0.15, 0.59]; RT: Pearson’s r = 0.432, *p* < 0.001, 95% CI = [0.20, 0.62]; Confidence: Pearson’s r = 0.639, *p* < 0.001, 95% CI = [0.46, 0.77]; Figure 7B) and MSDNet (Accuracy: Pearson’s r = 0.389, *p* = 0.002, 95% CI = [0.15, 0.59]; RT: Pearson’s r = 0.80, *p* < 0.0001, 95% CI = [0.69, 0.88]; Confidence: Pearson’s r = 0.853, *p* < 0.001, 95% CI = [0.77, 0.91]; Figure 7D), demonstrating that all three models predict better the data from individuals who behave more similarly to the rest of the group. However, BLNet, showed no significant correlation for accuracy predictions (Pearson’s r = −0.029, *p* = 0.828, 95% CI = [-0.28, 0.23]; Figure 7C) while exhibiting high correlations for RT (Pearson’s r = 0.831, *p* < 0.001, 95% CI = [0.73, 0.90]) and confidence (Pearson’s r = 0.809, *p* < 0.001, 95% CI = [0.70, 0.88]). All correlations were highest for RTNet compared to the other three networks. These analyses further support the notion that RTNet provides the best model of average human behavior among existing alternatives.

**Figure 7.**
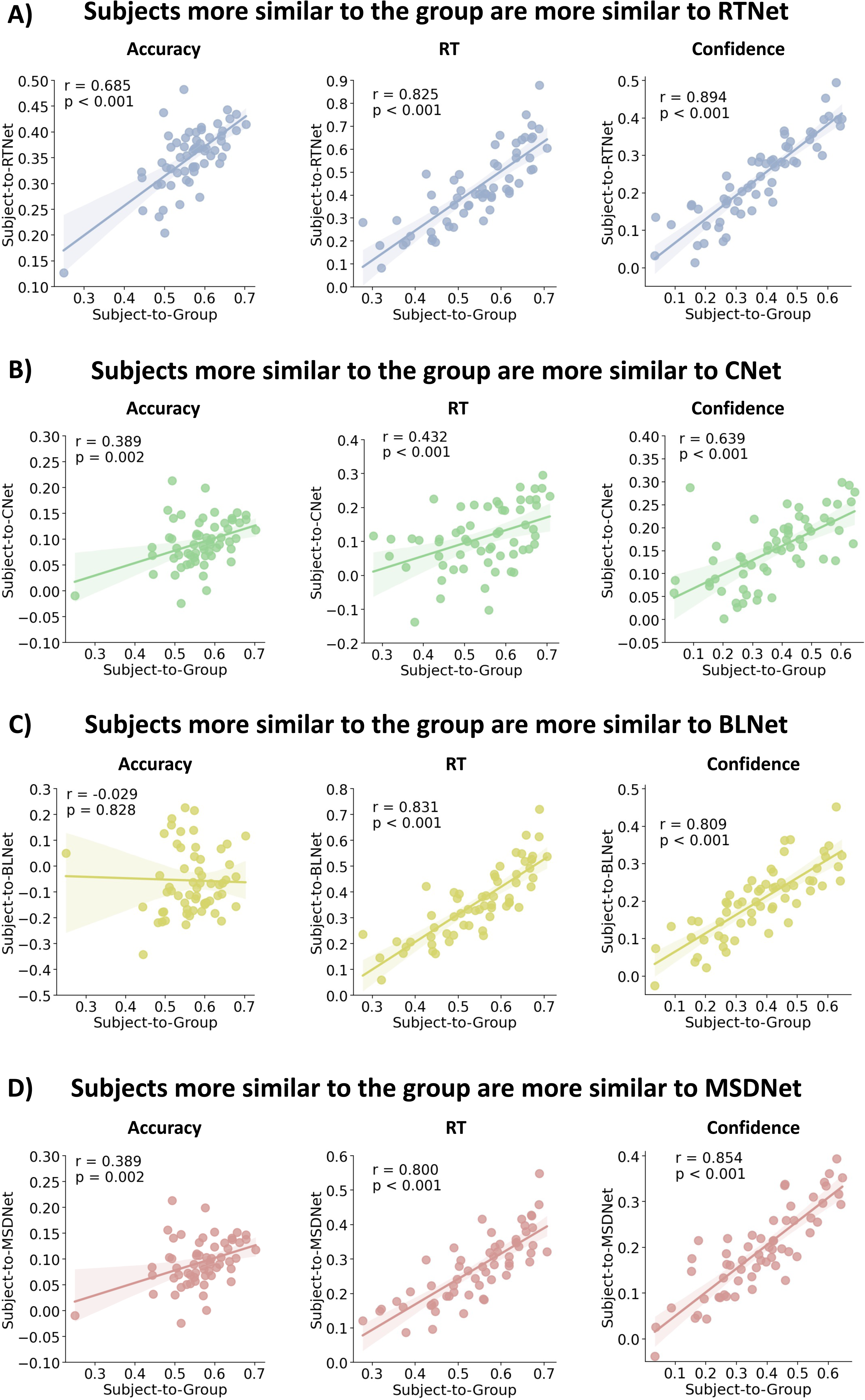
Humans who are more similar to the group average are also more similar to each model. (A) We observed a strong positive correlation between the subject-to-group and subject-to-RTNet similarity values for accuracy, RT, and confidence. This finding indicates that individual subjects whose behavior was more similar to the group average on per image basis were also more similar to the predictions made by RTNet. (B-D) Similar results were also observed for CNet, BLNet (except for accuracy correlations), and MSDNet, although these correlations tended to be lower than for RTNet. Dots represent individual subjects; lines depict best-fit regressions; shaded areas depict 95% confidence intervals around the regression estimate.

To understand better these results, we further examined who were the subjects whose accuracy, RT, and confidence was most similar to the group. We found that different subjects had the highest similarity to the group for RT compared to accuracy or confidence (Supplementary Figure 4A-C). Therefore, RTNet and other models did not simply provide good fit to specific subjects but instead provided good fits to different groups of subjects for different measures. Finally, the individuals closest to the group in their mean accuracy also tended to be those who had the highest task accuracy, suggesting that RTNet and the other models were better at predicting the image-by-image accuracy of subjects with higher task performance (Supplementary Figure 4D).

Given the variability in how similar individual subjects were to the group data, we also explored how well the models compared to the ability of individual subjects to predict the group data. Two-sided paired t-tests showed that RTNet outperformed individual human subjects in predicting the accuracy (t(59) = 4.076, *p* < 0.001, Cohen’s *d* = 0.526, 95% CI = [0.02, 0.06]), RT (t(59) = 16.174, *p* < 0.001, Cohen’s *d* = 2.088, 95% CI = [0.2, 0.25]), and confidence (t(59) = 10.927, *p* < 0.001, Cohen’s *d* = 1.411, 95% CI = [0.18, 0.26]) of the rest of group across all conditions (Figure 8). Impressively, RTNet outperformed every individual human subject in predicting the group RT and confidence results, as well as 73.3% of individual subjects in predicting accuracy. On the other hand, CNet was significantly worse than individual subjects in predicting group accuracy and RT but not confidence (Accuracy: t(59) = −42.425, *p* < 0.001, Cohen’s *d* = 5.477, 95% CI = [-0.4, −0.39]; RT: t(59) = −25.439, *p* < 0.001, Cohen’s *d* = 3.284, 95% CI = [-0.38, −0.32]; Confidence: t(59) = −0.361, *p* = 0.719, Cohen’s *d* = 0.047, 95% CI = [-0.05, −0.03]). BLNet was significantly worse than individual subjects in predicting group accuracy but predicted group RT and confidence better than individuals (Accuracy: t(59) = −68.395, *p* < 0.001, Cohen’s *d* = 8.830, 95% CI = [-0.67, −0.63]; RT: t(59) = 7.018, *p* < 0.001, Cohen’s *d* = 0.906, 95% CI = [0.07, 0.13]; Confidence: t(59) = 6.170, *p* < 0.001, Cohen’s *d* = 0.797, 95% CI = [0.08, 0.16]). Finally, MSDNet’s predictions of group accuracy and RT were significantly worse than those of human subjects but its predictions of group confidence were better than those of individual subjects (Accuracy: t(59) = y42.425, *p* < 0.001, Cohen’s *d* = 5.477, 95% CI = [-0.42, −0.39]; RT: t(59) = −4.019, *p* < 0.001, Cohen’s *d* = 0.519, 95% CI [-0.08, −0.03]; Confidence: t(59) = 5.266, *p* < 0.001, Cohen’s *d* = 0.68, 95% CI = [0.07, 0.15]). In sum, RTNet was the only network that outperformed most individual subjects in predicting all three measures of human performance (accuracy, RT, and confidence).

**Figure 8.**
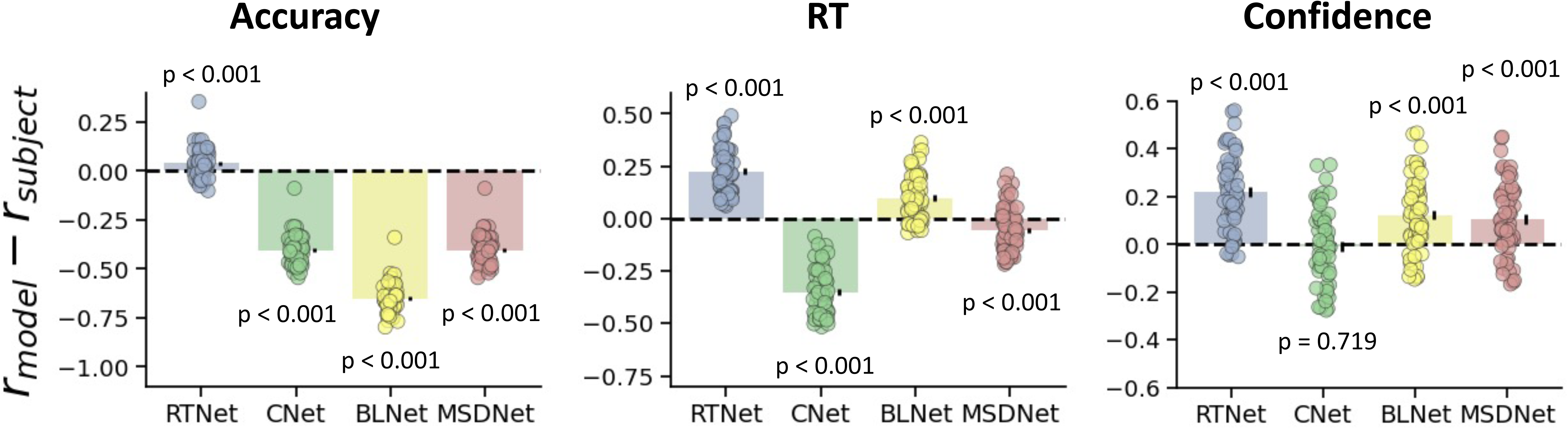
Comparison between individual subjects and the models in predicting the group data. RTNet significantly outperformed individual human subjects (n = 60) in predicting group accuracy, RT, and confidence. On the other hand, CNet, BLNet and MSDNet were worse than individual humans in predicting accuracy, and CNet and MSDNet were worse in predicting RT. We note that the effect sizes are very small for RTNet’s predictions of accuracy and MSDNet’s predictions of RT. However, the effect was sufficiently consistent across subjects to make these results statistically significant (RTNet outperformed 44/60 subjects in predicting accuracy and MSDNet did worse than 43/60 subjects in predicting RT). For all panels, dots represent individual subjects; error bars show SEM. The p-values are derived from two-sided one-sample t-tests.

## Discussion

There is considerable interest in using neural networks as models of human visual processing and behavior, but relatively little work has been done on testing the extent to which existing image-computable models reproduce the full range of behavioral signatures exhibited by humans. Here we show that the current state-of-the-art neural networks such as CNet, BLNet, and MSDNet diverge in several ways from human behavior. Further, we develop a new neural network, RTNet, that exhibits all critical features of human perceptual decision making, including effects on accuracy, RT, and confidence. Further, RTNet predicted well human group behavior for novel images and did so better than both CNet, BLNet, and MSDNet, as well as better than individual human subjects. Finally, individual humans who were more similar to the group were also more similar to RTNet. Overall, RTNet is a promising image-computable model of human accuracy, RT, and confidence.

### Relationship between RTNet and cognitive models of perceptual decision making

RTNet is the first neural network to exhibit all critical signatures of human perceptual decision making. This success, however, is hardly surprising given the strong conceptual similarity between RTNet and traditional cognitive models of decision-making that also exhibit the signatures of human behavior^24,26,40,52,54^. These models are often referred to as sequential sampling models where (usually noisy) evidence is accumulated over time until a threshold is reached. The most common sequential sampling models are diffusion models, which are typically only applied to 2-choice tasks where evidence in favor of one response alternative is also evidence against the other alternative^1,40^. Instead, RTNet is conceptually more similar to another subgroup of sequential sampling models called race models where each choice option has its own accumulation system and evidence for each choice is accumulated in parallel^42,55^.

Despite their conceptual similarity, RTNet has two important advantages over traditional cognitive models. Most importantly, RTNet is image-computable and can be applied to actual images, whereas traditional models cannot. As such, traditional models cannot replicate RTNet’s ability to make accurate predictions regarding human accuracy, RT, and confidence for individual unseen images. The second advantage stems from the inability of traditional cognitive models to naturally capture the relationships between the different choice options. Specifically, to maintain a low number of free parameters, cognitive models are often fit with the assumption that evidence accumulates at the same rate for all incorrect choice options (but accumulates faster for the correct choice)^56^. However, this assumption ignores the fact that some incorrect options may be more similar to the correct option and thus are more likely than other options to be chosen. While dependencies between the choices can easily be incorporated in cognitive models, that would result in a large number of free parameters that would make fitting to data difficult. Conversely, RTNet inherently learns all relationships between the choice options during the training of the Bayesian neural network that forms its core. RTNet still requires the fitting of the overall signal strength (which we accomplish by adjusting the noise level of the images fed to RTNet), but this single free parameter allows it to capture all choice option dependencies, something that traditional models cannot achieve.

### Performance differences between RTNet and other networks

RTNet outperformed all other networks we tested (CNet, BLNet, and MSDNet) in capturing the signatures of perceptual decision making. Specifically, while MSDNet and CNet show relatively weaker effects of task difficulty on RTs compared to humans, RTNet closely captures the observed magnitude of this effect. Further, RTNet is the only model that mimics the observed shape and skewness of RT distributions in response to SAT/difficulty manipulations. Finally, RTNet yielded the closest image-by-image predictions of human choice, RT, and confidence.

We speculate that RTNet’s ability to match observed patterns in human behavior, particularly RTs, is primarily due to its internal mechanisms being closer to the true mechanisms that give rise to RTs in humans. Specifically, RTNet’s core assumption that RTs are generated by a process of sequential sampling and evidence accumulation is inspired from a long tradition of cognitive modelling ^1,2^. In fact, these evidence accumulation models have been tested extensively against human data and are currently the best models of human RTs ^1,2^. On the other hand, models such as CNet, BLNet and MSDNet rely on mechanisms that, although can generate RTs, have not been as extensively validated by empirical tests and are therefore less likely to capture the true mechanisms that generate RTs in humans.

Nevertheless, another reason why CNet and MSDNet may struggle with generating human-like RTs is that the RTs generated by the models are constrained by the number of layers or residual blocks present in the networks. On the other hand, RTNet’s evidence accumulation mechanism allows flexible generation of RTs across a potentially very large number of steps, thus allowing the RTs to have higher resolution and sensitivity to experimental manipulations.

### Biological plausibility of neural network models of response time

Physiological recordings have uncovered several features of the processing in the human visual system that are relevant to judging the plausibility of the networks examined here. First, the conduction from one area to another in the visual cortex (roughly corresponding to different layers in neural networks) takes approximately 10 ms^57^, with signal from the photoreceptors reaching the top of the visual hierarchy in inferior temporal cortex in 70-100 ms^58^. Therefore, a single sweep from input to output in a purely feedforward network should result in decisions with RT less than a few hundred milliseconds even though human decisions can range from a hundred of milliseconds to a few seconds. Second, neurons in each layer of the visual cortex continue to fire action potentials for hundreds of milliseconds after the stimulus onset and receive strong recurrent input from later layers of processing^59^. Finally, neuronal processing is known to be noisy such that the same image input generates very different neuronal activations on different trials^37^.

MSDNet diverges from these known properties of the human visual cortex in several important ways. To generate meaningful RTs, MSDNet assumes that classification decisions are made after each layer of processing, though there is no evidence that decisions in the brain can be directly based on information in early visual cortex without further processing in subsequent layers. Moreover, because it assumes the existence of a single feedforward sweep through the network, it cannot naturally capture large RT variability between stimuli given the short latencies of processing between different layers. Finally, MSDNet does not incorporate any recurrent processing, capture the noisiness of the responses in the visual cortex, or replicate the long periods of activity of the neurons in each processing area. These properties strongly limit the biological plausibility of MSDNet.

In comparison, the dynamics of CNet are closer to those of biological neural networks. Indeed, several of CNet’s features – such as parallel and continuous processing of input, and transmission delays between layers – were directly inspired by biology. The transmission delays allow the network to mimic the processing latencies across cortical layers. These features were also found to account for differences in processing efficiency between images such that CNet produced more rapid responses for prototypical images with clear backgrounds compare to unusual or cluttered images. However, CNet includes several features that are not biologically plausible such as its lack of stochasticity of decisions and recurrent processing. Further, it remains unclear how its cascaded architecture could map onto brain areas^12^.

BLNet appears more biologically plausible than both MSDNet and CNet as it features recurrent visual processing. Lateral connections in RCNNs enable a layer’s activations from previous time steps to feed back into itself, which allows state dependence to naturally emerge in these networks, thus mimicking biological networks^67^. Additionally, RCNNs have been found to generate RTs that align closely with human RTs on a range of complex perceptual tasks involving scene categorization, perceptual grouping, and mental simulation^22^. These findings suggest further similarities in perceptual processing between humans and RCNNs. However, in spite of these advantages, RCNNs still lack certain features of biological networks such as stochasticity of responses.

It is possible to introduce stochasticity in CNet and MSDNet by feeding the outputs of the final softmax layer into a race model. However, such an architecture would imply that response stochasticity arises purely from noise in the decision stage. Although decision noise may exist in humans contributing to noisy motor responses, stochasticity in human responses is thought to predominantly arise from noisy inference^29^ or noisy sensory representations^60–62^. Therefore, CNNs with additional noise at the decision stage are less biologically plausible than RTNet, which includes noise in the evidence processing stage.

On the other hand, while also not capturing all properties of visual processing, RTNet appears more biologically plausible. First, it mimics the noisiness of neuronal responses for repeated presentations of the same stimulus. Second, through the process of evidence accumulation, RTNet naturally generates long-lasting neuronal activations. Third, the network’s output is inherently stochastic, unlike CNet, BLNet, MSDNet, or standard feedforward networks that are inherently deterministic. Finally, the accumulation process implemented in RTNet has been observed in multiple regions in the human parietal cortex, frontal cortex, and subcortical areas^63–66^. Nevertheless, one critical limitation of the biological plausibility of RTNet is its lack of recurrency. That being said, the question of how to train recurrent neural networks on static images remains open^53,58,67–69^. Further, while the core of RTNet does not include recurrency, the evidence accumulation system can be thought of as a recurrent network. In fact, several recent studies have demonstrated the advantages of combining a standard feedforward network with a recurrent network in performing a range of tasks and extrapolating to solve problems of greater complexity than they were trained on^70,71^. Future studies should explore how to introduce recurrence into RTNet’s mechanisms and whether such modifications can improve its predictions of human behavior.

### Using noisy weights to generate stochasticity in RTNet’s responses

One critical feature of RTNet is that its weights are noisy. Practically, there are many ways of generating noise in the weights. In early iterations of RTNet, we attempted to create variability by training a feedforward network and then adding the same amount of variability to each connection. This approach resulted in variability that was too small for some weights and too large for others^72^, often leading to no accuracy gains from the process of evidence accumulation. Indeed, a given amount of noise over a specific weight may not change the performance of a network at all, but the same disturbance over another weight may have destructive effects^73–75^. We therefore chose to obtain the weight variability by training a Bayesian neural network so that each weight has an appropriate amount of noise. In the future, it may be possible to use other methods for setting the noise level for each connection, but we are currently unaware of any method besides training a Bayesian neural network that can generate appropriate noise for each weight.

Another alternative to implementing noise in RTNet is to only add noise to the weights in the pre-readout layer (which can mimic noise in the decision process rather than in the sensory processing). As there are many different ways to implement stochasticity in the network, it is important for future studies to test how these differences in implementation affect the model’s performance.

RTNet is built such that every time evidence is sampled from a stimulus, the network’s weights change randomly (according to the BNN’s posterior weight distributions). These random moment-by-moment fluctuations in the network’s weights lead to noisy activations. However, in the brain, noisy activations in response to a stimulus are thought to arise from random fluctuations in neuronal activity itself. Therefore, it can be argued that a more biologically plausible implementation of RTNet would involve noise in unit activations rather than weights^76^. The main reason we chose to add noise in weights rather than activations is due to the practical ease of implementing BNNs that can naturally generate variability in networks. Mechanistically, however, there may be no meaningful distinction between noisy weights and noisy activations. Indeed, noisy weights lead to noisy activations, which mimic the randomness of neural responses.

### Limitations

One limitation of RTNet is that its mechanism for stopping the accumulation process is non-optimal. Following a large literature of race models in cognitive psychology^24,42,56^, RTNet makes a decision when any one choice option receives sufficient evidence to exceed a threshold. However, if another choice option has almost same amount of evidence, the observer has little ability to differentiate between the two choices and is essentially guessing between them. Previous research showed that guessing can be an appropriate behavior if the observer knows that the task is very difficult^77^ or if the observer has been deliberating for a long time^78^. However, in a race model, guessing can happen at any time point regardless of task difficulty. Nevertheless, human decisions are often suboptimal^79,80^, and therefore it is unclear as to whether this suboptimal decision-making mechanism should be seen as a drawback if the goal is to model human decision-making.

Another limitation of RTNet is that each sweep of the feedforward path is independent of the previous states, whereas the current state in the human brain is influenced by its previous states^67^. To address this limitation, the sampling process in RTNet can be modified such that the current state of the network depends on the previous states. For example, during testing, the connection weight at a specific moment can be made a function of its previous values, which would make the sequential samples dependent on each other. Additional studies are needed to investigate the effect of such state dependence on model performance.

### Conclusion

We developed a new neural network, RTNet, which exhibits the basic features of human perceptual decision making and predicts human accuracy, RT, and confidence on an image-by-image basis. The network provides a better model of human perceptual decisions than the current state-of-the-art networks for generating response times. RTNet thus represents an important step in the use of neural networks as models of human decisions.

## Methods

All subjects signed informed consent and were compensated for their participation. The protocol was approved by the Georgia Institute of Technology Institutional Review Board, protocol H15308. All methods were carried out in accordance with relevant guidelines and regulations.

### Behavioral experiment

#### Pre-registration

This study’s sample size, experiment design, included variables, hypothesis, and planned analyses were pre-registered on Open Science Framework (https://osf.io/kmraq) prior to any data being collected.

#### Subjects

Sixty-four subjects (31 female, age=18-32) with normal or corrected to normal vision were recruited. We had pre-registered the collection of only 40 subjects, but due to less time restrictions than we had anticipated, and to further increase the statistical power, we collected data from more subjects.

#### Stimulus, task, and procedure

Subjects performed a digit discrimination task where they reported their perceived digit followed by rating their decision confidence. Each trial began with subjects fixating on a small white cross for 500-1000 ms, followed by a presentation of the stimulus for 300 ms (**Error! Reference source not found.**). The stimulus was a digit between 1 and 8 (the digits 0 and 9 were excluded) superimposed on a noisy background. Subjects’ task was to report the perceived digit using a computer keyboard by placing four fingers of their left hand on numbers 1-4 and placing four fingers of their right hand on numbers 5-8. This setup allowed subjects to respond without looking at the keyboard, thus providing less noisy response times. Following their categorization response, subjects reported their decision confidence on a 4-point scale (where 1 corresponds to the lowest confidence and 4 corresponds to the highest confidence). There was no deadline on the response or confidence rating.

The experiment included manipulations of speed-accuracy trade off and task difficulty. Speed-accuracy trade off was manipulated by asking subjects to emphasize either the speed or accuracy of their responses. To facilitate proper responding, we organized the experiment into alternating blocks of speed and accuracy focus. Task difficulty was manipulated by adding different levels of uniform noise to the stimuli. Specifically, “easy” stimuli included average uniform noise of 0.25 (range = 0-0.5), whereas “difficult” stimuli included average uniform noise of 0.4 (range = 0-0.8). To add the noise, the pixel values were first transformed to be between 0 and 1 and random numbers drawn from the corresponding noise distributions were added separately to each pixel. We scaled the resulting image to be between 0 and 1 again, and finally converted the image to a uint8 format (scaled between 0 and 255). The noise levels were chosen based on the pilot testing to produce two different performance levels. Easy and difficult images were randomly interleaved.

The task stimuli were selected from a publicly available handwritten digits (MNIST) dataset^32^. This dataset contains 60,000 training images and 10,000 testing images. Since the training images were used to train the models in this study, we randomly selected images from MNIST test set to include in our experiment. This ensures that the selected images for the experiment are novel both for the human subjects and for the trained models. We randomly selected 480 images for the experiment (120 for each condition). The MNIST dataset images are of size 28 x 28 pixels which appeared overly small on the computer screens we were using. Therefore, before adding noise, the selected images were first resized to 84 x 84 pixels (using MATLAB’s *imresize* function), and they were padded with the background color of MNIST images to size 256 x 256 pixels (visual angle = 6.06°).

The experiment started with three blocks of training each containing 50 trials. The first block contained images from the MNIST dataset without any noise. This was done to familiarize the subjects with the experiment. The next two blocks were used to introduce the speed-accuracy trade off by asking subjects to focus on accuracy in the first block and on speed in the second. The noise level of the stimuli in these two training blocks was same as in the main experiment (i.e., 0.25 and 0.40 for the easy and difficult stimuli, respectively). During the whole training session, the experimenter was standing beside the subject quietly and was available to answer any questions. None of the images used in the training session was used in the main experiment.

Once the subject confirmed that he or she understands the task, the experimenter left the room and subjects completed the main experiment that consisted of 960 trials organized in four runs each containing four blocks of 60 trials. Each block consisted of a single speed-accuracy trade off condition, and each run included exactly two “accuracy focus” and two “speed focus” conditions in a randomized order. At the beginning of each block, subjects were given the name of the condition for that block (“accuracy focus” or “speed focus”) and asked to adjust their responding policy accordingly. In each block, we pseudo-randomly interleaved trials from the two difficulty levels such that each was presented exactly 30 times. All 480 images were shown to subjects in first two runs and the procedure was repeated with a new random ordering of the stimuli in the last two runs. All images were same for all subjects, and each image was assigned only to one specific condition.

#### Apparatus

The experiment was designed in MATLAB 2020b environment using Psychtoolbox 3^81^. The stimuli were presented on a 21.5-inch Dell P2217H monitor (1920 x 1080 pixel resolution, 60 Hz refresh rate). Subjects were seated 60 cm away from the screen and provided their responses using a keyboard.

### Behavioral analyses

We followed the data analyses steps outlined in our preregistration. All analyses were performed in Python (version 3.10.11) using Google Colab (version 2.0). We first excluded subjects who did not follow sufficiently well the speed/accuracy instructions by not providing faster average RT in the “speed focus” compared to the “accuracy focus” condition. This resulted in removing two subjects (out of 64). We preregistered the exclusion of subjects with floor or ceiling effects on accuracy but no subject met the criteria for exclusion. However, following our preregistration, we excluded two subjects because they showed ceiling effects for confidence. Note that our preregistration document called for excluding subjects who provided average confidence of more than 3.7 but because this would have resulted in excluding a much larger number of subjects than we had anticipated, we only excluded subjects whose average confidence was above 3.85. Therefore, 60 subjects were used in all subsequent analyses.

We additionally excluded individual trials with extreme RT values using preregistered criteria based on Tukey’s interquartile criterion. Specifically, for each subject, we computed the 25^th^ and 75^th^ percentiles of the RT distributions in each condition. We then removed all RTs with values more than 1.5 times the interquartile range such that if *Q*1 is the RT value at the 25^th^ percentile and *Q*3 is the RT value at the 75^th^ percentile, we removed values smaller than *Q*1 − 1.5 × (*Q*3 − *Q*1) and larger than *Q*3 + 1.5 × (*Q*3 − *Q*1). This step resulted in removing an average of 5.46% of total trials (range of 1.35-8.22% for each subject).

Once these preprocessing steps were completed, we computed average accuracy, RT, confidence, and skewness of the RT distributions separately for each condition. The skewness was computed separately for each individual subject’s RT distribution as 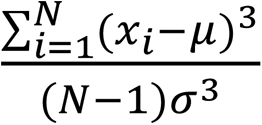 where *μ* and *σ* are the mean and standard deviation of the sample distribution, respectively. We also computed average RT and average confidence scores for error and correct trials across subjects to examine how RT and confidence change as a function of response accuracy. Finally, for visualization purposes, we plotted RT distributions for one subject in Figure 4C. The RT distributions were generated using kernel density estimation (KDE), which approximates the underlying probability density function that generated the data by smoothing the observations with a Gaussian kernel^82^. The KDE plots were created using Seaborn’s KDE plot with a smoothing bandwidth of 1.2^83^.

### RTNet

#### Network architecture

The RTNet model consists of two main modules (A). The first module is a Bayesian neural network (BNN) which makes predictions regarding an image. BNNs are a type of artificial neural network built by introducing stochastic components into the network to simulate multiple possible models with their associated probability distribution^84^. The main difference between a BNN and standard feedforward neural network is that in BNN the weights are distributions instead of point estimates. A random sample from these distributions results in a unique feedforward network. This random sampling enables variability in the output of the network, which in turn can be fed into an accumulation process that drives a decision. The second module of our model consists of exactly such accumulation of the evidence produced on each step by the first module. At each processing step, the output of the network (in the form of activations of the final layer) was accumulated towards a pre-defined threshold. Evidence for each choice option was accumulated separately from the rest, similar to a race model^24^.

The accumulation process continues until the total amount of accumulated evidence for one of the alternatives reaches a predefined threshold. The alternative for which the threshold was reached then becomes the response of the model. The response time produced by RTNet is simply the number of samples used to reach the decision threshold. The confidence of the model was obtained by taking the difference in evidence scores between the chosen response and the second-best choice.

#### Implementation

We implemented RTNet using the AlexNet architecture, which has eight layers with learnable parameters^33^. The AlexNet architecture consists of five convolutional layers with a combination of max pooling followed by three fully connected layers. We chose to implement RTNet within a relatively large-scale CNN such as AlexNet (rather than a shallow network which may have also been able to learn to classify the MNIST dataset) because our goal was to eventually compare our model to others such as CNet and MSDNet, which are generally based on larger CNNs and work on multiple existing datasets. Additionally, difficulties associated with training Bayesian neural networks limited us to relatively small network structures (rather than VGG or ResNet models). We found the AlexNet architecture to be a reasonable compromise in this trade off between model complexity and ease of training BNNs. RTNet was implemented in PyTorch^85^ while Bayesian networks were implemented using Pyro^86^, which is a probabilistic programming library built on PyTorch^85^.

#### Training the BNN module of RTNet

BNNs are probabilistic models that incorporate uncertainty into their weights and biases, rather than treating them as point estimates. Consider a training dataset, *x*, for which we must predict the class labels, *y*. In traditional neural networks, the predicted class label, [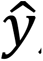 is a function of network’s weights, *w*, and these weights are tuned in order to optimize the correspondence between the predicted (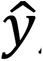) and true class labels (*y*). In BNNs, however, weights are modelled as probability distributions instead of point estimates. Following the rules of Bayesian inference, one can infer the posterior distribution of these weights (*w*) using the formula 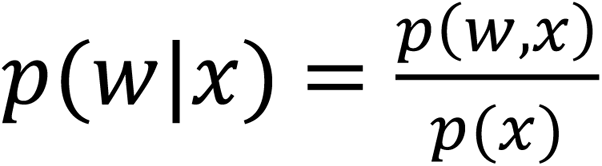.However, this computation is intractable for large networks since it involves computing the marginal likelihood of the data *p*(*x*) across all possible configurations of weights. Therefore, computing this posterior distribution is typically done using a method of approximation called variational inference. A stand-in distribution, *q*(*w*), is specified to approximate the posterior and its parameters are tuned to maximize the similarity between the two distributions. The similarity between the distributions is quantified by the information theoretical measure called Kullback-Liebler (KL) divergence:

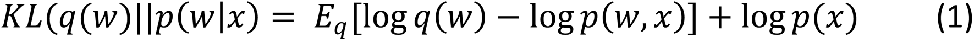

Although *KL*(*q*(*w*)||*p*(*w*|*x*), cannot be directly computed since *p*(*x*) is intractable, one can side-step this computation by defining a surrogate objective function called the evidence lower bound (ELBO) function as:

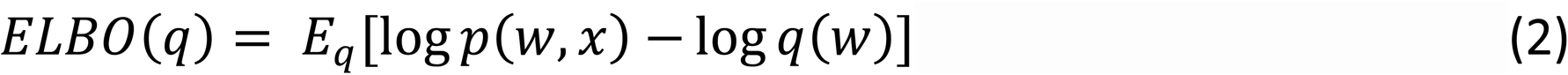

where both *p*(*w*, *x*) and *q*(*w*) are tractable, and due to their negative relationship, maximizing *ELBO*(*q*) thus results in the minimization of *KL*(*q*(*w*)||*p*(*w*|*x*), allowing one to approximate the true posterior distribution of the network’s weights.

We trained the network to achieve classification accuracy higher than 97% on the MNIST test set. We trained the BNN module of RTNet for a total of 15 epochs with a batch size of 500. We used the Evidence lower bound (ELBO) loss function^87^ and Adam^88^ for optimization with a learning rate of 0.001, and the default values for weight decay and epsilon (weight decay = 0; ε = 10^-8^). To ensure that each BNN performs greater than 97% on MNIST test set, we followed a specific rule for each model instance. When testing an image with the BNN module of RTNet, we sampled 10 times from the posterior distributions learned during the training and thus obtained 10 unique responses for each image. The response with highest frequency among 10 responses was chosen as the final decision of the BNN module. Note that there were no RTs generated at this step since we only implemented the BNN module of RTNet and generated a set of responses that would allow us to evaluate how well the BNN’s posterior distributions had been trained. These trained BNN models were later used to generate variable activations for the evidence accumulation process that resulted in RTs.

We resized the MNIST images to the standard input size to AlexNet model architecture (227 x 227 pixels). We also normalized the input images to have a mean of 0.1307 and standard deviation of 0.3081, which is a standard procedure when using AlexNet for classification of the ImageNet dataset^89^. We trained sixty instances of RTNet using the above procedure but with different weight initializations for each network instance. We used a different combination of mean and standard deviation (SD) values for each of the 60 instances to maximize differences in network initializations. Specifically, different network instances of RTNet were initialized such that all means of the weights and biases were set to a value between 0.1 and 1.2 with 0.1 increments, and all SDs of weights and biases were set to a value ranging from 1 to 5 with increments of 1 (for a total of 12 × 5 = 60 instances).

#### Generating RTNet’s responses from the evidence accumulation module

Sequential sampling models belong to a class of cognitive models which assume that observers make decisions by repeated sampling and accumulation of noisy evidence until a threshold is reached ^1,2^. In these models, RT reflects the number of sampling steps required to reach the threshold. RTNet utilizes this evidence accumulation mechanism to generate RTs. In order to generate noisy evidence, we used the probability distribution of weights in the BNNs to randomly sample one unique feedforward network at each time step. At each time step, *t*, the presented image results in a feedforward sweep of the sampled network and generates a set of activations (*a*_t_) where *a*_t_ = [*a*_1,t_, *a*_2,t_ … *a*_8,t_] are the values obtained in the last layer after the softmax function has been applied. Each unit in the output layer corresponds to the activation for one of the eight choice options and for each choice, the evidence obtained at the current step is added to the sum of evidence collected from all previous steps. Thus, a running total of accumulated evidence is maintained such that 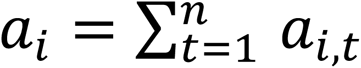 where *n* refers to the total number of steps over which evidence has been accumulate and *i* ∈ [1,8] refers to the response option. When the total evidence in favor of any of the options exceeds a pre-defined threshold *k*, the corresponding response option is chosen such that the network’s response, *r* = argmax(*a*_1_ *a*_2_… *a*_8_) at the time step when max (*a*_1_, *a*_2_ … *a*_8_) ≥ *k*.

What are the properties of evidence accumulation? Everything else being equal, decisions that are based on fewer evidence samples are more likely to be influenced by chance fluctuations in evidence that favor incorrect decisions. On the other hand, when the model is allowed to accumulate evidence over a longer period, these such random variations are more likely to cancel out, thus increasing the likelihood of a correct response. In turn, because a longer period of accumulation leads, on average, to stronger evidence, this directly results in higher confidence.

### CNet

#### Network architecture

The parallel cascaded network (CNet) builds upon the architecture of residual networks (ResNet) by utilizing skip connections to introduce propagation delays during input processing (Figure 1B). At each processing step, all units in all layers are updated parallelly. However, due to the propagation delays introduced by each residual block, simpler perceptual features get transmitted faster across blocks. For instance, at the first time-step, only the first residual block receives input and model predictions at this step are based only on the computations of the first residual block. At the second time step, all the other layers receive partial input from the first block. Even though the model prediction at this point will be based on computations from all blocks, only the first block will have received complete input and achieved stable output. The other blocks will only contain partial updates from the lower block and therefore their output will not be stable. In general, a residual block, *t*, takes (*t* − 1) time steps to receive complete and stable input. At any point during processing, the network can generate a prediction since all the residual blocks contribute to the computations. However, if the time step (*t*) is less than the number of residual blocks, the responses will be based on unstable representations in the higher blocks. Due to this architecture, the network’s responses are subject to a trade off between speed and complexity of processing. Decision time is indicated by the processing step at which the decision was made, and decision confidence is derived from the softmax value in the final layer, at the time of decision. The softmax values are obtained by transforming the activation scores (*z*) of all nodes in the output layer according to the function: 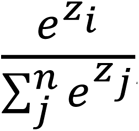, where *i* refers to the node whose output is being transformed and *n* refers to the number of nodes in the output layer (which is equal to the number of classes).

#### Implementation

CNet was implemented using the architecture of ResNet-18^9^ since it requires networks with skip connections. ResNet-18 architecture consists of 17 convolutional layers, where 16 of these layers are embedded within eight residual blocks (skip connections), followed by a final fully connected layer with softmax activation to generate the decision. The network was implemented in PyTorch^85^.

#### Network training

We trained CNet using the same procedure that was used by the original authors since their training protocol was found yielded the best network behavior and performance. The network achieved an accuracy > 97% with 12 training epochs and a batch size of 500. The models were trained on a temporal-difference (TD) learning procedure along with cross-entropy loss. In the original publication, TD learning was found to perform better than softmax-based cross-entropy loss in encouraging correct responses to emerge faster. The loss function was optimized using an initial learning rate of 0.01, weight decay of 0.005 and a momentum of 0.9. The images were normalized to a mean of 0.1307 and standard deviation of 0.3081. We trained sixty instances of CNet using the above procedure but using a different random seed for initializing the network’s weights to allow individual differences in network’s learning.

### BLNet

#### Network architecture

BLNet is a recurrent convolutional neural network (RCNN) consisting of a standard feedforward CNN with recurrent connections that connect each layer to itself^10^ (Figure 1C). A final readout layer computes the network’s output at each time step via a softmax function. Time steps are defined as the number of feedforward sweeps of the network that have occurred until the time at which the readout is evaluated. At each time step, a given layer receives input from two sources – the feedforward input from the previous convolutional layer and recurrent input from itself in the form of activations from the previous time step. The readout is evaluated at each time step such that if it exceeds a pre-defined threshold, the network generates a response. The response corresponds to the choice that generates the highest softmax value and the time step at which the response was made indicates the decision time. The softmax value associated with the choice at the time of decision indicates decision confidence. The network’s ability to trade off speed and accuracy comes from the fact that higher softmax thresholds require more feedforward and recurrent computations, which effectively results in a deeper network being unrolled across time which, in turn, leads to both higher RT and higher accuracy.

#### Implementation

BLNet was implemented as a custom-built network consisting of seven convolutional layers of increasing size and a final readout layer, as defined by the original authors^10^. Each layer consists of two sets of weights – the bottom up weights that transform the input from the previous layer and the lateral weights that act on recurrent input that the layer receives from itself. The readout layer is a fully connected layer with softmax activation to generate the decision. The network was unrolled across time for eight time steps. The network was implemented using TensorFlow.

#### Network training

We were able to achieve test accuracy > 97% with only three epochs with a batch size of 32 and a sparse categorical cross-entropy loss function^90^. Adam^88^ was used for optimization with a learning rate of 0.001. For testing, the response at the final time-step was taken as the network’s decision. We resized the MNIST images to the standard input size of 128×128 defined for the network. We trained sixty instances of BLNet using the above procedure but using a different random seed for initializing the network’s weights to allow individual differences in network’s learning.

#### Testing

Unlike the other networks, BLNet exhibited overall accuracy that was about 5% greater for the 120 images used in the easy, speed-focus condition compared to the 120 images used in the easy, accuracy-focus condition. This resulted in a lack of the expected accuracy difference between these two conditions when BLNet was run on all images (Supplementary Figure 5). On further investigation, we found that for each condition, the image set contained a small subset of images for which the network showed chance-level performance (12.5%). The image set for the easy, accuracy-focus condition contained more such images than the image set for the easy, speed-focus condition, explaining the observed accuracy differences. Therefore, when testing BLNet on the effects reported in Figure 4, we excluded this subset of images for all conditions (10 out of 480 images). This exclusion led to BLNet showing the expected speed-accuracy trade off (Figure 4A,B).

### MSDNet

#### Network architecture

MSDNet has an architecture similar to a standard feedforward neural network but with early-exit classifiers after each of its layers (D). At each output layer, the evidence for each choice is computed using a softmax function and if the evidence for any alternative exceeds a predefined value the network stops processing and immediately produces a response. The layer at which the response was made is indicative of the decision time, and the softmax value at that layer is indicative of decision confidence^90,91^.

#### Implementation

We implemented MSDNet using the AlexNet architecture, which has eight layers with learnable parameters^33^. The AlexNet architecture consists of five convolutional layers with a combination of max pooling followed by three fully connected layers. In addition to the standard AlexNet structure, we incorporated additional readout layers located right after each layer of processing. The feature map size of all these readout layers were set to the number of classes. The network was implemented in PyTorch^85^.

#### Network training

Due to MSDNet’s deterministic nature, only three epochs with a batch size of 500, were enough to achieve test accuracy of more than 97% with the same batch size and a weighted cumulative loss function^90^. Adam^88^ was used for optimization with a learning rate of 0.001. For testing, the response of the last output layer was taken as the network’s decision. If a network did not achieve accuracy greater than 97%, we started the training over with the same initial values. Since MSDNet is also built on the architecture of AlexNet, we resized the MNIST images to the standard input size for AlexNet and normalized the images to have a mean of 0.1307 and standard deviation of 0.3081. To make the initializations of MSDNet as similar as possible to the initializations of RTNet, for each RTNet instance, we set the initial values for the weights and biases of the MSDNet instance by randomly sampling from the Gaussian distribution used in the corresponding RTNet initialization.

### Choosing parameters that allow the models to mimic human accuracy

Because the goal of our study was to examine whether the models exhibit the signatures of human perceptual decision making, we matched the accuracy of the models across the four experimental conditions to the average accuracy in the human data. For all models, this was achieved by adjusting the noise level in the images (separately for the “easy” and “difficult” images) and the threshold parameter (separately for the speed and accuracy conditions). Lower noise levels lead to higher accuracy, whereas higher threshold parameters lead to longer processing and response times (and also contribute to higher accuracy levels).

Parameter values were adjusted using a coarse search followed by a fine search. In the coarse search for RTNet, we varied the amplitude of uniform noise from 1 to 10 with increments of 1 (where the noise amplitude refers to the length of the interval over which the noise values are generated), and the threshold value from 2 to 12 with increments of 2. The results were closest to the human accuracy levels when the noise was in the range 2-3 for easy images and 4-5 for difficult images, and the threshold was set to 2-4 for the speed focus condition and 6-8 for the accuracy focus condition. We then conducted a fine search near those values by changing the noise level from 2 to 5 with 0.1 increments and changing the threshold values from 2 to 8 with 0.5 increments. The closest match to human accuracy was achieved for noise levels of 2.1 and 4.1 for easy and difficult images, respectively, and a threshold value of 3 for the speed condition and 6 for the accuracy condition. With these threshold and noise parameters, the evidence accumulation process in RTNet executed 6.5 sampling steps on average, although the distributions were wide such that the actual steps varied from 1 to 35. However, the number of processing steps depended on the experimental manipulation with the number of steps increasing for both difficult images and with stress on accuracy over speed (the average number of steps observed for each condition correspond to the height of the bars for RTNet in Figure 4B).

We used a similar procedure to tune the parameters of CNet, BLNet and MSDNet. Note that the threshold value for CNet is the softmax evidence at the output layer. The coarse search was performed using threshold values between 0.5 and 0.9 with increments of 0.04. The results were closest to the human accuracy levels when the threshold was in range 0.79-0.83 for the speed focus condition, and 0.86-0.9 for the accuracy focus condition. We then performed a fine search in these ranges by incrementing the threshold by steps of 0.01. The closest match to human accuracy was achieved for a threshold value of 0.83 for the speed condition and 0.9 for the accuracy condition. For noise levels, the best match to human accuracy was obtained when the noise levels were set to 1.42 and 1.83 for easy and difficult images, respectively. For BLNet, like CNet, the threshold value is the softmax evidence at the output layer. The coarse search was performed using threshold values between 0.1 and 0.95 with increments of 0.2. The results were closest to the human accuracy levels when the threshold was in range 0.4-0.5 for the speed focus condition, and 0.9-0.95 for the accuracy focus condition. We then performed a fine search in these ranges by incrementing the threshold by steps of 0.05. The closest match to human accuracy was achieved for a threshold value of 0.4 for the speed condition and 0.95 for the accuracy condition. For noise levels, the best match to human accuracy was obtained when the noise levels were set to 0.55 and 1.2 for easy and difficult images, respectively.

The threshold value for MSDNet is the softmax evidence at each early exit. The coarse search was performed using the threshold values between 0.5 and 0.95 with increments of 0.05. The results were closest to the human accuracy levels when the threshold was in range 0.55-0.65 for the speed focus condition, and 0.8-0.9 for the accuracy focus condition. We then performed a fine search in these ranges by incrementing the threshold by steps of 0.01. The closest match to human accuracy was achieved for a threshold value of 0.58 for the speed condition and 0.82 for the accuracy condition. For finding the optimal noise levels, the best match was obtained when the noise levels were set to 1.9 and 3.0 for easy and difficult images, respectively.

Although we tried to closely match each network’s accuracy with that of humans for each condition, our ability to do this was limited by the fact that a given SAT threshold must predict accuracies for both the easy and difficult conditions and a given noise level must predict accuracies for both the SAT conditions. Therefore, we obtained parameters estimates that resulted in closely (but not exactly) matched accuracies.

## Data availability

Behavioral data have been made publicly available at: https://osf.io/akwty.

## Code availability

All codes and trained models are publicly available at: https://osf.io/akwty.

## Acknowledgments

This work was supported by the National Institute of Health (award: R01MH119189) and Office of Naval Research (award: N00014-20-1-2622), both awarded to D.R. The funders had no role in study design, data collection and analysis, decision to publish or preparation of the manuscript We thank Sashank Varma and Paul Verhaeghen for helpful suggestions about the analyses, as well as Ana Shin and Himanaga Sahithi Pandi for assistance with data collection.

## Author Contributions

F.R. and M.S. performed research and analyzed the data; F.R. collected the data and wrote the first draft of the paper; M.S. and D.R. edited the paper; All authors designed research.

## Competing interests

The authors declare no competing interests.

## Notes

### Competing Interest Statement

The authors have declared no competing interest.

### Summary of Updates

RCNN is added to the group of models that we compared RTNet against them.

