## supplementary materials for "RTNet neural network exhibits the signatures of human perceptual decision making"

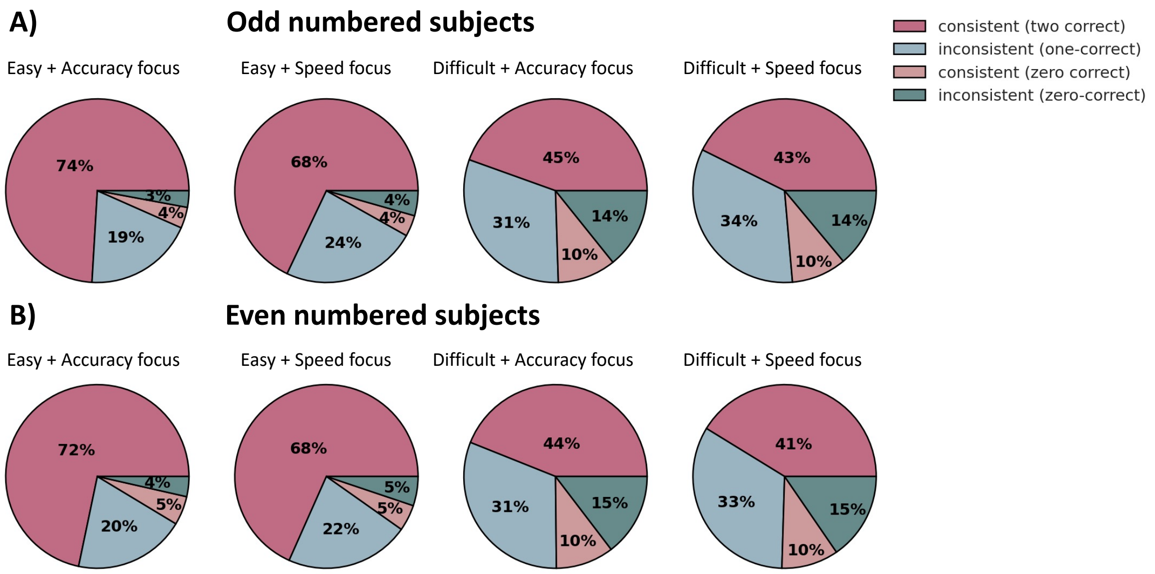


**Supplementary Figure 1. Testing the robustness of our measure of stochasticity.** We computed stochasticity as the proportion of trials where the subject (n = 60)/network's responses were inconsistent between the two repeats of the same image. However, it is possible that having only two repeats per image does not allow us to estimate a sufficiently reliable measure of stochasticity. Therefore, to test whether our estimates of stochasticity were reliable, we computed this measure separately for odd and even numbered subjects (which is similar to assessing split-half reliability of a measure). The proportions of inconsistent responses were not significantly different between the two halves of the sample (Two sided Wilcoxon's signed rank test: $Z(59) = 3850.5$, $p = 0.716$, rank-biserial correlation (effect size) = -0.03). Further, the two estimates never diverged by more than 2%, confirming that our estimates were reliable and robust. In the legend, "consistent (two correct)" refers to instances when the correct responses was given for both presentations of a given image, "consistent (zero correct)" refers to instances when the same incorrect choice was made both times, "inconsistent (one correct)" refers to instances when only one of the choices was correct, "inconsistent (zero correct)" refers to instances where different incorrect choices were made each time.

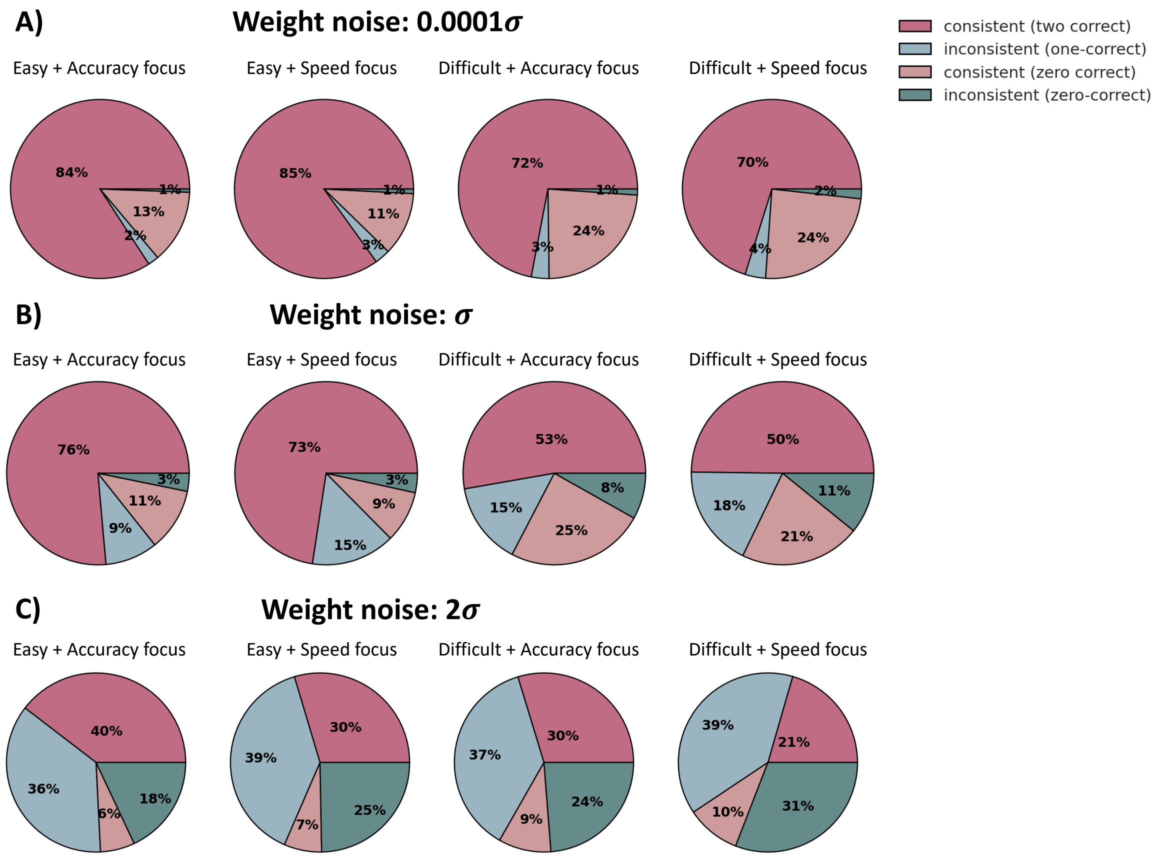


**Supplementary Figure 2. Effect of changing weight noise on decision stochasticity in RTNet.** RTNet exhibits stochastic decision-making with stochasticity increasing with task difficulty and speed stress. Here, we tested whether the stochasticity of the network's responses (model instances = 60) can be manipulated by changing the variance of the network's weight distributions. We manipulated the noise in the weights by multiplying the variance of the weight distributions in the BNN by a factor of A) 0.0001, B) 1 and C) 2 and observed that the network's stochasticity increased with the variance of its weights. The overall stochasticity (proportion of inconsistent responses for the same image) across the four experimental conditions increased from 4% to 61% as the variance of the weights was scaled from 0.001 to 2. We also observed an overall decrease in accuracy across all experimental conditions with increase in weight noise. In the legend, "consistent (two correct)" refers to instances when the correct responses was given for both presentations of a given image, "consistent (zero correct)" refers to instances when the same incorrect choice was made both times, "inconsistent (one correct)" refers to instances when only one of the choices was correct, "inconsistent (zero correct)" refers to instances where different incorrect choices were made each time.

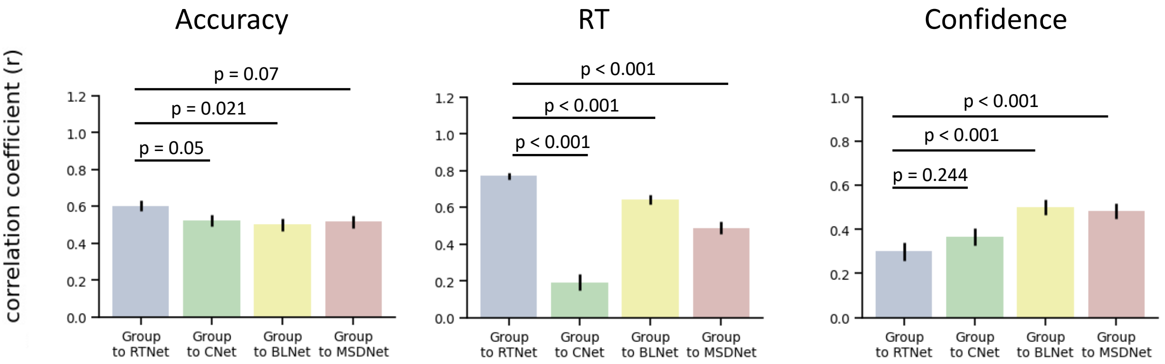


**Supplementary Figure 3. Image-by-image correlation between average human data and average model predictions**. We assessed the correlations between human accuracy, RT, and confidence and the corresponding quantities predicted by the models across individual images (n = 480) by pooling all four experimental conditions. In each case the model prediction was derived by averaging the behavior of the 60 instances of the networks for a given image. We found that all models predicted average human behavior on individual images well for accuracy (RTNet: Pearson's r = 0.603, 95% CI = [0.54, 0.66]; CNet: Pearson's r = 0.523, 95% CI = [0.45, 0.59]; BLNet: Pearson's r = 0.499, 95% CI = [0.43, 0.56], MSDNet: Pearson's r = 0.516, 95% CI = [0.45, 0.58]; all *p's* < 0.001), RT (RTNet: Pearson's r = 0.770, 95% CI = [0.73, 0.8]; CNet: Pearson's r = 0.191, 95% CI = [0.1, 0.28]; BLNet: Pearson's r = 0.643, 95% CI = [0.59, 0.69]; MSDNet: Pearson's r = 0.488, 95% CI = [0.42, 0.55]; all *p's* < 0.001), and confidence (RTNet: Pearson's r = 0.299, 95% CI = [0.22, 0.38]; CNet: Pearson's r = 0.366, 95% CI = [0.29, 0.44]; BLNet: Pearson's r = 0.501, 95% CI = [0.43, 0.57]; MSDNet: Pearson's r = 0.483, 95% CI = [0.41, 0.55]; all *p's* < 0.001). Using the Fisher's r-to-z transformation and two-sided tests, we found that RTNet produced the highest correlation with accuracy compared to other networks. However, these differences were only significant for BLNet (z(479) = 2.315, *p* = 0.021) and marginally significant for CNet (z(479) = 1.813, *p* = 0.070) and MSDNet (1.961, *p* = 0.050). For RT, the correlations between human subjects and model predictions were significantly higher for RTNet compared to CNet (z(479) = 12.771, *p* < 0.001), BLNet (z(479) = 3.970, *p* < 0.001) and MSDNet (z(470) = 7.519, *p* < 0.001). For confidence, on the other hand, CNet, BLNet and MSDNet were found to have higher correlations compared to RTNet although these differences were only significant for BLNet and MSDNet (CNet: z(479) = -1.164, *p* = 0.244; BLNet: z(479) = -3.741, *p* < 0.001; MSDNet: z(479) = -3.374, *p* < 0.001). Error bars show SEM derived parametrically from the Pearson's correlation coefficient ($r$) and the sample size ($n$ = 480 images) as $\sqrt{\frac{1-r^2}{n-2}}$.

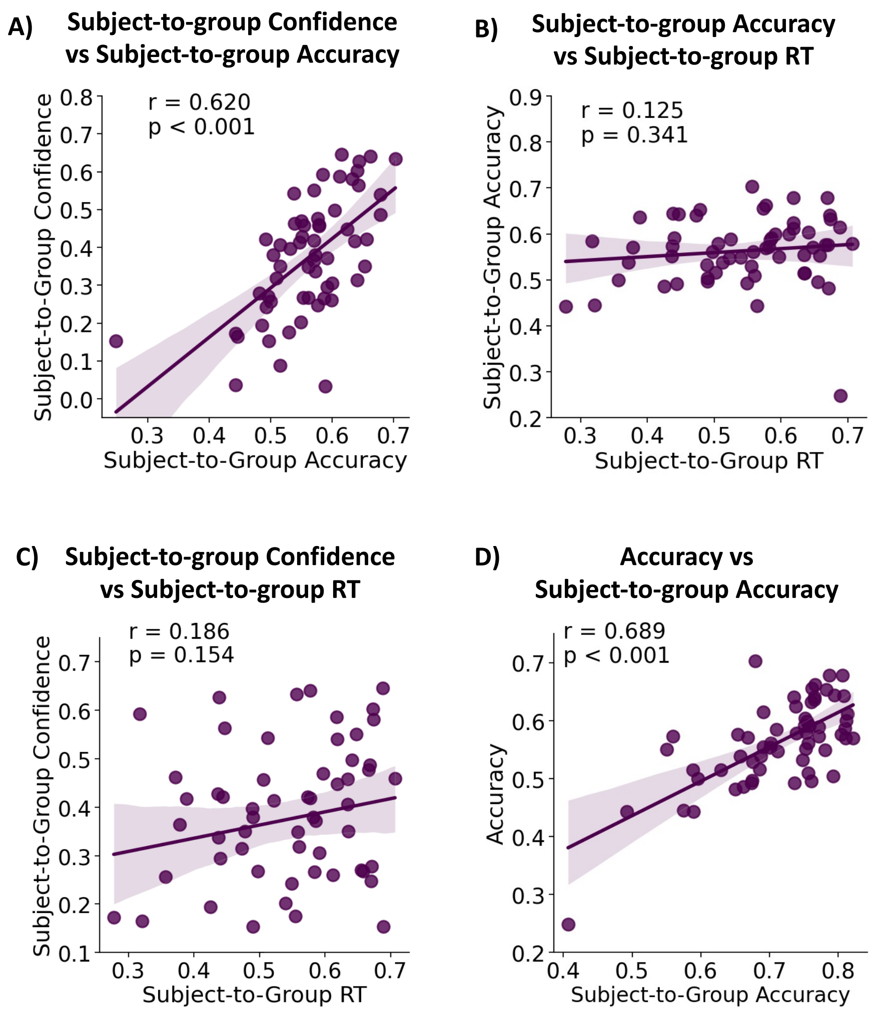


**Supplementary Figure 4. Assessing how subjects' similarity to the group varies between different measures (accuracy, RT and confidence) and with task performance.** We found that RTNet best predicted those subjects (n = 60) who were closest to average human behavior. However, it is unclear whether a subject who is similar to the group for one measure (e.g., accuracy) would also be similar to the group for the other measures (e.g., RT and confidence). Therefore, here we test whether the same or different subjects show the highest similarity to the group between the three measures. A) We observed a high correlation between subject-to-group similarity in confidence and subject-to-group similarity in accuracy (Pearson's r = 0.620, p < 0.001, 95% CI = [0.43, 0.75]). These results suggest that subjects who are closest to the group's accuracy are likely to be the same subjects who are closest to group mean confidence. B,C) However, unlike the relationship between accuracy and confidence, there was no strong association between RT and accuracy (Pearson's r = 0.125, p = 0.341, 95% CI = [-0.13, 0.37]) or RT and confidence (Pearson's r = 0.186, p = 0.154, 95% CI = [-0.07, 0.42]). Therefore, the subjects who were closest to the group's RTs were mostly different from those who were closest to the group's accuracy and confidence. In other words, RTNet does not simply provide a good match to specific subjects, but matches different subjects for different measures based on how similar subjects are to the group in that specific measure. D) We tested how the subject's similarity to the group relates to their task performance. We found a high correlation between task accuracy and subject-to-group accuracy (Pearson's r = 0. 689, p < 0.001, 95% CI = [0.53, 0.8]), implying that subjects who were closest to the group accuracy and thus best predicted by RTNet were those who showed the highest task performance. Dots represent individual subjects; lines depict best-fit regressions; shaded areas depict 95% confidence intervals around the regression estimate.

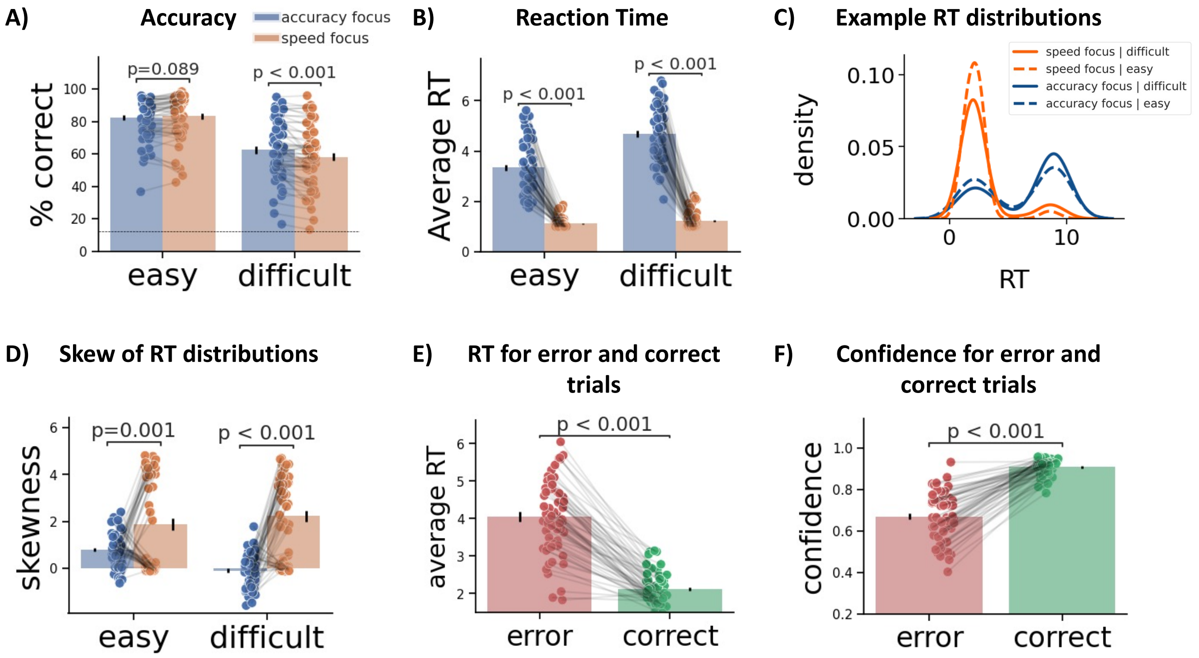


**Supplementary Figure 5. Behavioral effects shown by BLNet on the full experimental dataset**. On the original MNIST test images used for the experiment, BLNet showed no difference in accuracy between the speed- and accuracy-focus conditions for easy images. However, for the difficult condition, there was a significant decrease in accuracy for the speed-focus condition. When we examined the networks' responses for images within each experimental condition separately, BLNet showed an overall higher accuracy (by ~5%) for the images in the easy, speed-focus condition compared to the easy, accuracy-focus condition. This accuracy difference likely masked the accuracy drop resulting from speed pressure for the easy images. Therefore, we excluded a small subset of images for which the network showed chance-level performance (accuracy < 12.5%; 10 out of 480 images). Here we report the results for BLNet (n = 60 model instances) without exclusions. (A) Accuracy for each of the four conditions, showing no accuracy difference for the two easy conditions without image exclusions. (B-F) All other results remained similar to the results after image exclusions reported in Figure 4. For all panels, dots represent individual subjects; error bars show SEM. The p-values are derived from two-sided Wilcoxon's signed rank tests (for mean RT comparisons) and two-sided paired t-tests (for all other measures).
